# Spatially resolved proteomics of the stomatal lineage: polarity complexes for cell divisions and stomatal pores

**DOI:** 10.1101/2023.11.03.564551

**Authors:** Eva-Sophie Wallner, Andrea Mair, Dominik Handler, Claire McWhite, Shou-Ling Xu, Liam Dolan, Dominique C. Bergmann

## Abstract

Cell polarity is used to guide asymmetric divisions and create morphologically diverse cells. We find that two oppositely oriented cortical polarity domains present during the asymmetric divisions in the Arabidopsis stomatal lineage are reconfigured into polar domains marking ventral (pore-forming) and outward facing domains of maturing stomatal guard cells. Proteins that define these opposing polarity domains were used as baits in miniTurboID-based proximity labeling. Among differentially enriched proteins we find kinases, putative microtubule-interacting proteins, polar SOSEKIs with their effector ANGUSTIFOLIA, and using AI-facilitated protein structure prediction models, we identify their potential interaction interfaces. Functional and localization analysis of polarity protein OPL2 and its newly discovered partners suggest a positive interaction with mitotic microtubules and a potential role in cytokinesis. This combination of cutting-edge proteomics and structural modeling with live cell imaging provides insights into how polarity is rewired in different cell types and cell cycle stages.

## Introduction

Polarity proteins orient cell divisions, promote cell fate diversity, and guide the construction of specialized cell morphologies during the development of complex multicellular organisms [1, 2]. Especially in plants, where cells cannot migrate, the activities polarity proteins regulate are essential to generating organized and well-functioning tissues and organs [3]. In eudicot plants like *Arabidopsis thaliana*, the epidermal lineage that produces stomata has become a paradigm to dissect the interplay between cell fate, cell division, and cell polarity during tissue patterning. The multipotent stomatal lineage produces, as one product, stomata, which are cellular valves comprised of two kidney-shaped sister guard cells that flank a pore. By regulating the aperture of this pore, stomata mediate gas exchange between the plant and the surrounding environment. The stomatal lineage features an early stem-cell like proliferative stage where oriented asymmetric cell divisions (ACDs) generate meristemoids (eventual guard cell precursors) and stomatal lineage ground cells (SLGCs) that can become pavement cells or re-enter a stem-cell stage by later ACDs. Differentiation of stomata follows a fate conversion from meristemoid to guard mother cell (GMC), the symmetric division of the of GMC to form guard cells, and subsequent growth and morphological changes to generate a pore between the guard cells.

Proteins responsible for size and fate asymmetries during cell divisions include the polarly localized proteins BASL [4], BASL’s interaction partners in the BREVIS RADIX (BRX) family [5], and POLAR [6, 7]. BASL, BRX family and POLAR are intrinsically disordered and each serves as a scaffold for a different intracellular signaling cascade [5, 8] [6]. Nuclear migration and destabilization of microtubules within the BASL-BRX domain orients the division plane away from the polar domain to ensure asymmetry of the division [9, 10]. After ACD, the larger SLGC daughter therefore inherits the BASL-POLAR-BRX polarity domain including the scaffolded effectors, whereas the smaller meristemoid daughter does not, leading to the two daughters acquiring different fates [5, 6, 8]. Opposite to this BASL-POLAR-BRX polar domain is a second domain marked by OCTOPUS-LIKE (OPL) proteins, especially OPL2 [11]. OPLs do not have a clear role in orienting ACDs but do have a role in daughter cell fate. OPLs are segregated to the smaller meristemoid daughter of an ACD and function to promote continued proliferation (“stem-ness”) in meristemoids [11].

Many plant cell types are polarized and use common polarity scaffolds. For example, it was in developing phloem that opposite polarization of OPL2 and BRXL2 homologues was first observed. OPL2 and OCTOPUS (OPS) polarize to the shootward side of developing phloem cells to promote differentiation into sieve elements [12, 13], whereas BRX polarizes to the rootward side where it fine-tunes phloem differentiation [14, 15]. Despite the parallels in their polar localization in phloem and the stomatal lineage, the functions of, and interactions between, OPL and BRXL families differ depending on the cellular context [11]. We hypothesize that seemingly contradicting roles of OPL and BRXL proteins in the stomatal and phloem lineages have their origin in distinct protein partners of OPLs and BRXLs. BRXL and OPL (and BASL and POLAR) proteins are largely unstructured [16, 17] with intrinsically disorganized regions (IDRs) that are a common feature of scaffolding proteins [18]. These properties make functional analysis, protein extraction and *in vitro* protein assays challenging as the stability of such proteins often largely depends on their microenvironment and presence of their interaction partners.

Here, using TurboID-based proximity labeling, we generate local proteomes of the two opposing polarity domains marked by OPL2 and BRXL2 and identify stomatal-lineage specific potential interaction partners of OPL2. We also reveal novel polarity patterns of OPL2-BRXL2 in relation to microtubules in symmetrically dividing guard mother cells and discover a role of OPLs in the initiation of the stomatal pore. Taking advantage of new AI-facilitated protein structure predictions, we identify potential protein-protein interfaces between OPL2 and the polarity protein SOSEKI3 (SOK3) [19], DYNEIN LIGHT CHAIN 1 (DLC1) [20] and multiple subunits of CASEIN KINASE II (CKII) [21]. Functional and localization analysis of OPL2 and its newly discovered partners suggest a positive interaction with mitotic microtubules and a role in aiding cytokinesis, as OPL2, SOK3 and DLC1 co-localize at specific regions of the phragmoplast. These cutting-edge proteomics and structural models, combined with live imaging, provide insights into how polarity is rewired in different cell types and cell cycle stages and an inventory of spatially restricted plasma-membrane associated proteins able to participate in specialized plant cell divisions and morphogenesis.

## Results

### OPL marks dynamic cortical polarity domains during the symmetric guard mother cell division

We previously showed that OPLs mark a novel polarity domain in asymmetrically dividing stomatal lineage cells that is inherited by the meristemoid daughter, and that OPLs collectively promote continued asymmetric divisions in these cells [11]. We noticed, however, that peak OPL2 accumulation (from reporter *OPL2p:OPL2-VENUS)* occurred later during formation of the paired stomatal guard cells (Figure 1A-B’). We therefore carefully tracked OPL2 dynamics from meristemoids (Figure 1C-C’) through the symmetric cell division of guard mother cells (GMCs) (Figure 1D-I’). In young GMCs, OPL2-VENUS occupied two opposing polar patches that marked the long axis of the GMC (Figure 1D-D’). These patches coincide with regions of cell wall thickening described in [22]. As cytokinesis commences, OPL2-VENUS also localizes to the growing cell plate (Figure 1E-E’) until the cell plate fuses with the two polar OPL2-patches (Figure 1F-F’). Newly divided GMCs show bright OPL2-VENUS signal at both sides of the newly formed cell wall (Figure 1G-G’), and OPL2 persists as the pore forms between the two guard cells (GCs) (Figure 1H-H’). Mature stomata were devoid of any OPL2-VENUS signal, indicating that OPL2 may play a role during stomata development, but not in the function of mature stomata (Figure 1I-I’).

**Figure 1:**
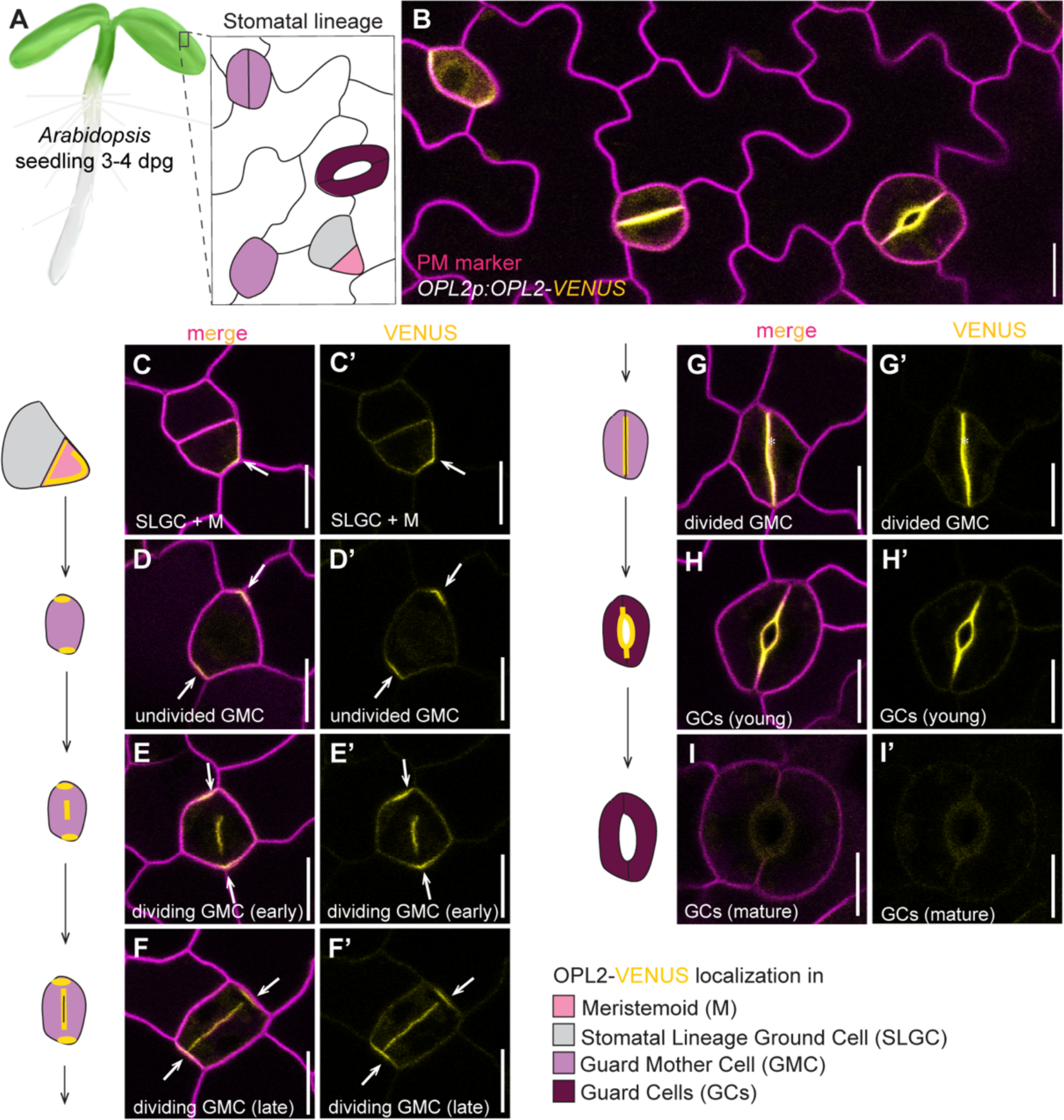
OPL2 polarizes in two patches marking the long axis of symmetrically dividing GMCs and extends along the pore-forming walls in young stomata. **(A)** Schematic representation of an *Arabidopsis thaliana* seedlings at 3-4 days post germination (dpg) with cotyledons displaying a variety of stomatal lineage stages within the epidermis (see key for identities). **(B-I)** Confocal images of cotyledon epidermis expressing *OPL2p:OPL2-VENUS* (yellow) and plasma membrane (PM) marker *35Sp:PIP2A-RFP* (magenta), C’-I’ are presented without PM channel. To the left of each panel pair C-I is a schematic representation of the OPL2 (yellow) distribution. (B) Overview of cell type-specific OPL2-VENUS polarization in GMCs. (C-C’) OPL2-VENUS is inherited by the smaller meristemoid after ACD [11]. (D-D’) OPL2-VENUS accumulates in two polar patches marking the long axis of the GMC just before symmetric cell division (SCD). (E-E’) OPL2-VENUS also accumulates at the growing cell plate during cell division, eventually fusing with the polar patches (F-F’). (G-G’) After SCD is completed, the polar OPL2-VENUS patches disappear, and signal is restricted to PM regions adjacent to the ventral walls separating the young GCs. (H-H’) OPL2-VENUS signal retracts from the edges of the new cell walls and accumulates around the forming pore in maturing GCs. (I-I’) Mature stomata lack OPL2-signal. White arrows point to polar OPL2-VENUS domains. Scale bars represent 10 µm. See also Figure S1

The emergence of two polar OPL2-VENUS patches before cell plate formation is provocative because GMCs deviate from the “shortest wall” division pattern typical of plants, where divisions are oriented along the shortest distance between mother cell walls [23]. The placement of the division plane and new wall is prefigured by the preprophase band of microtubules, and we hypothesized that the OPL2 patches could serve as guideposts to reorient the preprophase band along the long cell axis in GMCs. To enable high-resolution imaging of OPL2 and preprophase bands during the GMC division, we created lines co-expressing microtubule marker *TMMp:mCherry-TUA5* [24] and OPL2 driven by strong and GMC/GC-specific *FAMA* promoter [25] (*FAMAp:OPL2-VENUS)*. Indeed, OPL2-VENUS polar patch formation in GMCs coincided with preprophase band microtubule formation (Figure S1A-A’). Polar OPL2-VENUS patches stably marked the long axis throughout mitosis (Figure S1B-B’) and phragmoplast formation (Figure S1C-D’) until the cell plate fuses with the mother cell walls (Figure S1E-E’). In maturing stomatal guard cells, the origin of radial microtubule arrays overlaps with OPL2 at the ventral (“inner”) cell walls (Figure S1E-E’ and S1F-F’) [22, 26].

The co-localization of OPL2 and microtubules is especially relevant because of the recent discovery that the oppositely polarized BASL/BRXL2 complex orients division planes during ACDs by creating a zone that excludes stable preprophase band and interphase cortical microtubules [24]. We tested whether OPL2 and BRXL2 maintained their opposite polar localization and relationship to microtubules during symmetric GMC divisions; indeed, they do (Figure 2A-G) with OPL2 occupying the ventral and BRXL2 the dorsal (“outer”) domains of the newly divided cell pair (Figure 2C-C’). Co-monitoring a microtubule marker (*TMMp:mCherry-TUA5)* revealed similar relationships between polarity proteins and microtubule abundance in meristemoids (Figure 2D-E) and in dividing GMCs (Figure 2F-G) with the BRXL2 domain devoid of microtubules and the OPL2 domain coinciding with the preprophase band, the phragmoplast and cortical microtubules (Figure 2E-F, 2H and S1).

**Figure 2:**
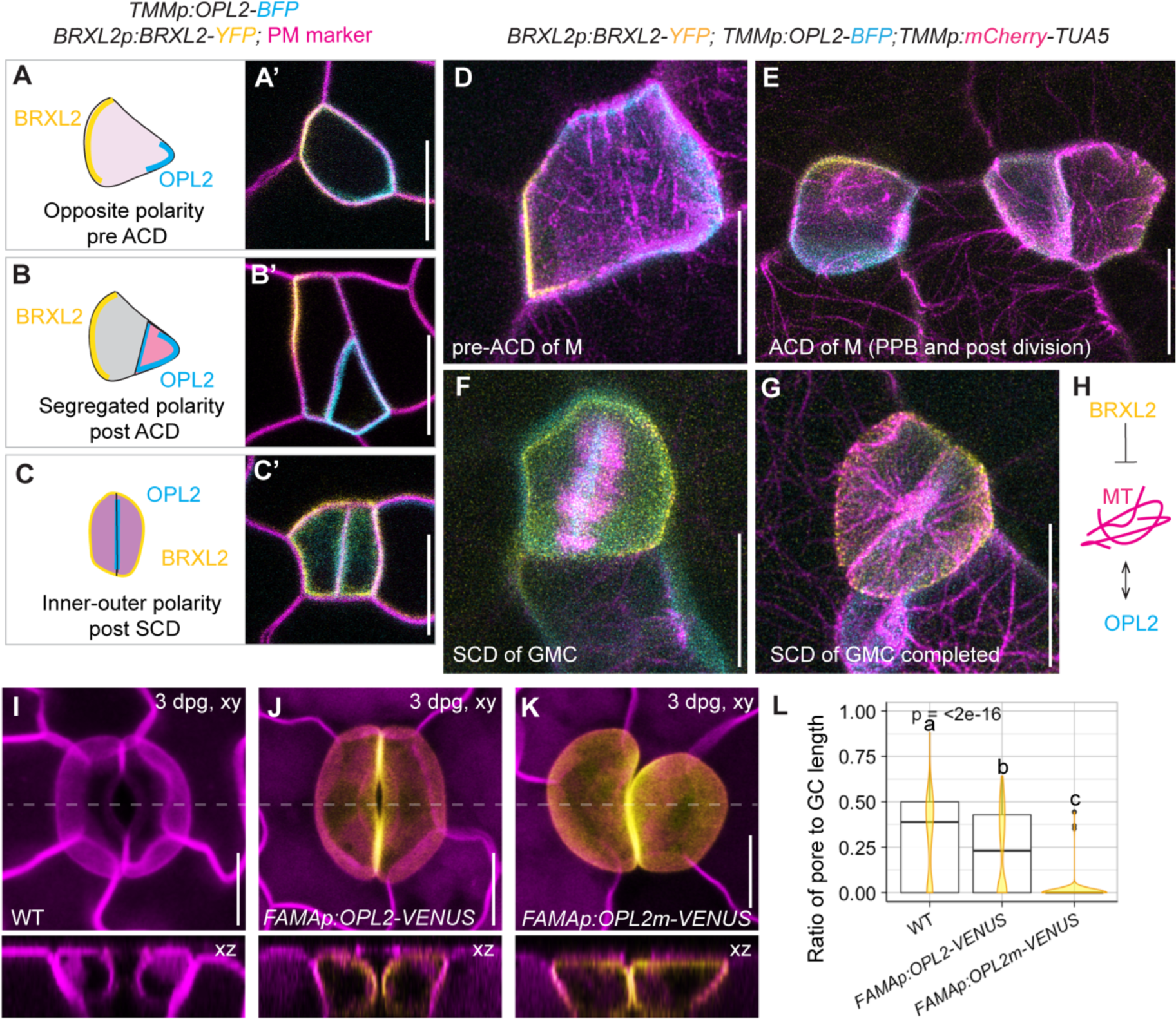
OPL2 and BRXL2 define opposite polarity domains with different relationships to microtubules. **(A-C)** Cartoons and confocal images (A’-C’) of the bipolar domains marked by *BRXL2p:BRXL2-YFP* (yellow) and *TMMp:OPL2-BFP* (cyan) plasma membranes are in magenta. In meristemoids, BRXL2 and OPL2 localize at opposing PM domains (A) pre-asymmetric cell division (ACD) and are (B) segregated to distinct daughter cells post-ACD. (C) In symmetrically dividing GMCs, they localize to ventral (‘inner’) (OPL2) and outer (BRXL2) domains. Scale bars represent 10 µm. **(D-G)** Meristemoids (E) and GMCs (F) expressing *BRXL2p:BRXL2-YFP* (yellow), *TMMp:OPL2-BFP* (blue) and stomatal lineage specific microtubule maker *TMMp:mCherry-TUA5* (magenta) show that BRXL2 avoids, but OPL2-BFP colocalizes with microtubules. Scale bars represent 10 µm. **(H)** Schematic representation of the relationships between polarity proteins BRXL2 (yellow), OPL2 (cyan) and microtubules (magenta). While BRXL2 destabilizes microtubules [24], OPL2 co-localizes and potentially interacts with microtubules (this paper). **(I-K)** 3D reconstructions of mature stomata expressing a PM marker *ML1p:mCherry-RCI2A* (magenta, I), *FAMAp:OPL2-VENUS* (J), *FAMAp:OPL2m-VENUS* (yellow, K) are depicted. Optical cross sections through xz are indicated by a dashed line in. *FAMAp*-expressed OPL2m-VENUS (J) inhibits stomatal maturation and pore formation when compared to wild type (I) and OPL2-VENUS (J). Scale bars represent 10 µm. **(L)** GC lengths and corresponding pore lengths were quantified in abaxial cotyledons of wild type, FAMAp:OPL2-VENUS and FAMAp:OPL2m-VENUS expressing lines. Measurements were taken from 291×291 µm regions flanking the midvein of 10 independent cotyledons per genotype. Within these regions, n = 119-148 stomata per genotype were measured and the ratio of pore length to total GC lengths was calculated and depicted as box plots with overlayed violin plots in yellow. A pore to GC length ratio of 0 indicates that no pore was detected. Groups represent statistical differences based on adjusted p-values < 0.05 determined by a Kruskal-Wallis multiple comparison followed by a post hoc Dunn test with p-values adjusted by the Holm method. See also Figure S2

Co-localization between OPL2 and microtubules could result from several different relationships; OPL2 could require microtubules for its localization, microtubules could be organized by OPL2, or both could respond to an upstream signal. Depolymerization of microtubules by short-term (30 minute) oryzalin treatments impaired preprophase band formation without disrupting the polar OPL2-VENUS patches marking the presumptive division plane along the long axis of the GMC (Figure S1G-G’). Likewise, attachment of OPL2-VENUS to the growing cell plate was stable and independent of microtubules (Figure S1H-H’) as was OPL2-VENUS localization to the ventral GC walls (Figure S1I-I’ and S1J-J’). These data suggest that OPL2 reads specific plasma membrane-localized marks that coincide with, but are largely independent of, microtubule structures before, during and after mitosis.

To investigate possible functions of OPLs during GMC divisions, we revisited tools that provided insight into OPL functions in meristemoid fate [11]. We generated lines expressing wild type OPL2-VENUS or the more stable OPL2^S295K;E296K^-VENUS (hereafter referred to OPL2m-VENUS) [11, 17] under control of the *OPL2* or *FAMA* promoters (Figure 2I-K and S2). While GMC divisions appeared to be normally oriented and cytokinesis was complete, pore formation of *FAMAp:OPL2-VENUS* was delayed relative to wild type and was almost completely absent in *FAMAp:OPL2m-VENUS* lines (Figure 2J-L). Over time, divided GMCs in *FAMAp:OPL2m-VENUS* lines expanded, but their shared (pore-forming) wall did not, resulting in large, deformed GCs that failed to form a stoma (Figure 2K and S2D-E). Timely expression of OPL2 may therefore be critical to coordinate pore formation with stomatal morphogenesis.

### Proximity labeling of polar domains in stomatal lineage cells reveals scaffolding partners and regulators of OPL2

Stomatal lineage polarity proteins are unstructured with few clues to their function encoded in their primary sequence. To identify molecular features and mechanisms underlying the distinct behaviors and functions of the BRXL/BASL/POLAR and OPL polar domains during asymmetric divisions and stomatal formation, we sought proteins that colocalize and preferentially interact with each polarity complex. We adapted a biotin-based proximity labeling method used previously for nuclear proteins in the stomatal lineage [27]. Additional optimization included choices of ligase, promoters, baits and controls. Compared to nuclear localized transcription factors, polarity proteins like OPL2 relocate quickly and frequently [11], so we made fusions between our bait proteins and mini-Turbo-ID (mTb), which is a small biotin ligase that covalently links biotin to proteins in close proximity of the bait under incubation conditions suitable for plant growth (Figure 3A) [27, 28]. We chose BRXL2 and OPL2 as the core baits because we find both to be exclusively plasma membrane associated, with BRXL2 doing so via palmitoylation [5]. Neither has been subjected to protein interaction assays, but their homologues BRX and OPL2, which display opposite polar localization in developing phloem, have some candidate interactors which can serve as references for evaluating our experiment [14, 15, 29].

**Figure 3:**
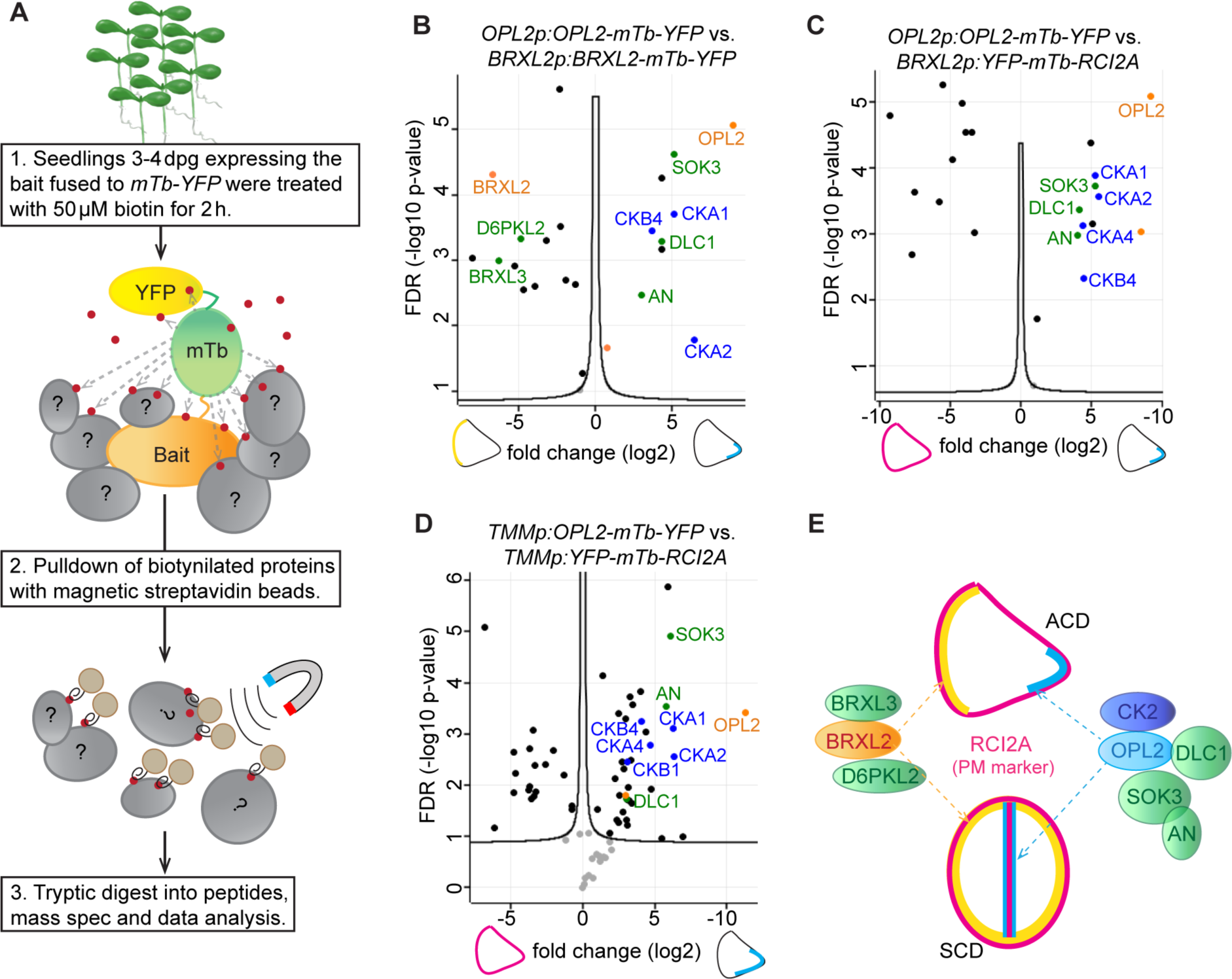
Proximity-labeling based proteomics of cortical polar domains in stomatal lineage cells identifies DLC1, SOK3, AN and CK2 peptides as enriched in OPL2 domains. **(A)** Schematic representation of the proximity labeling workflow to identify potential OPL2 interaction partners. 4 day-old seedlings expressing mTb-YFP tagged OPL2 (blue), BRXL2 (yellow) or RCI2A (pink) were used as bait (details in methods). (**B-D**) Volcano plots depict significantly (FDR = 0.5, S0 = 1) enriched proteins identified between independent triplicated samples expressing: (B) *OPL2p:OPL2-mTb-YFP* vs. *BRXL2p:BRXL2-mTb-YFP*, (C) *OPL2p:OPL2-mTb-YFP* vs. *BRXL2p:YFP-mTb-RCI2A* and (D) stomatal lineage specific *TMMp:OPL2-mTb-YFP* vs. *TMMp:YFP-mTb-RCI2A*. Proteins enriched in all OPL2 samples are DLC1, SOK3, AN (highlighted in green), casein kinase 2 (CK2) subunits (blue). Self-labeling of either BRXL2, OPL2 or mTb-YFP is depicted in orange. Pre-filtering removed wild type background (t-test: FDR = 0.5, S0 = 3) and only significantly enriched proteins with peptides present in all the three sample replicates were used for analysis (see methods). (**E**) Visual summary of novel potential protein-protein interaction partners in the BRXL2 and OPL2 polarity crescents, respectively. Based on expression domains of promoters used to drive baits, protein proximity could be in asymmetrically dividing meristemoids or symmetrically dividing GMCs See also Figure S3 and S4 and Table S1

We fused the mTb biotin ligase and a YFP tag to the C-termini of BRXL2 and OPL2. BRXL2-mTb-YFP under its native promoter (*BRXL2p:BRXL2-mTb-YFP*) rescued the stomatal phenotypes of the *brx-q* mutant (Figure S3A-C) and was localized to its expected polarity domain and subsequently inherited by the SLGC daughter after asymmetric division (Figure S3C-D). OPL2-mTb-YFP polarized as expected to the polar domain inherited by the meristemoid and to the newly formed ventral wall of young GCs (Figure S3E-F and H). Because we wanted to make comparisons among polarity domains, we also expressed OPL2-mTb-YFP under three promoters, *BRXL2p, OPL2p* and *TMMp* (Figure S3E, F and H), that represent, respectively: (1) a direct comparison with *BRXL2* expression, (2) the native *OPL2* expression and (3) strong but exclusively stomatal lineage expression. As controls for non-polarized plasma membrane association we also made mTb-YFP fusions to RARE-COLD-INDUCIBLE 2A (RCI2A) [30] driven by *BRXL2p* (Figure S3G) or *TMMp* (Figure S3I). RCI2A is a protein that we and others consistently use as PM marker [11, 31, 32] because RCI2A fusions to fluorescent proteins are uniformly distributed along the whole plasma membrane [33].

We treated 4-day-old seedlings expressing the respective mTb-YFP constructs (Figure 3A) with 50 µM biotin for 2 h (see methods) and a small portion of each sample was used for crude extracts and Western blotting to determine labeling efficiency (Figure S4). In addition to the endogenously biotinylated proteins [27], all mTb-expressing lines showed additional protein bands when compared to the wild type control (Figure S4). After this quality control step, we proceeded with protein extraction, pulldown of biotinylated proteins with streptavidin beads and finally mass spectrometry analysis. Data processing with MaxQuant [34] and Perseus [35] allowed us to extract the significantly enriched proteins per sample after filtering out the wild type background (see methods). Using volcano plots, we compared different sample groups (Figure 3B-D and S3J-K).

We first opted to make stringent comparisons between BRXL2-mTb-YFP and OPL2-mTb-YFP samples (Figure 3B and S3J), which grant us insights into the proteomes of two oppositely polarized protein domains in both stomatal and protophloem cells [11, 36]. 24 proteins were called differentially enriched between BRXL2-mTb-YFP and OPL2-mTb-YFP samples when expressed under their own promoters (Figure 3B and Table S1) and 33 proteins were differentially enriched when *BRXL2p* was used as a common promoter (Figure S3J and Table S1). Both comparisons yielded largely the same top hits. BRXL2-mT-YFP showed enrichment of its homolog BRXL3, which co-localizes with BRXL2 and is strongly expressed in the stomatal lineage [5] and D6PKL2, which was identified by immunoprecipitation as a BRX interactor in developing phloem cells [14].

The OPL2-mTb-YFP samples showed enrichment of the polarity protein SOK3 and its effector ANGUSTIFOLIA (AN) [37], DLC1, a poorly characterized microtubule-associated protein abundant in all plants (Cao et al., 2017) and multiple regulatory (CKB) and catalytic (CKA) subunits of protein kinase CKII [21]. CKII is ubiquitous and multifunctional, participating in several eukaryotic developmental pathways [38, 39]. The CKII target recognition motif is known and is present in OPL2 [17, 40]. Additional sample comparisons, such as OPL2 vs. the RCI2A non-polar PM line, result in enrichment of these same OPL2-sample dependent proteins (Figure 3C, S3K and Table S1).

Because the *OPL2* and *BRXL2* promoters drive expression in the root apical meristem [11], it was possible that OPL2 interaction partners were root-specific. We therefore used the *TMM* promoter to drive stomatal lineage specific expression of OPL2-mTb-YFP and the YFP-mTb-RCI2A plasma membrane control. Comparing these two lines yielded even more significantly enriched proteins, likely due to the fact that the *TMM* promoter is considerably stronger than the *OPL2* or *BRXL2* promoters [11], but among the top candidates enriched in *TMMp:OPL2-mTb-YFP* expressing lines we found again SOK3, AN, DLC1 and CKA and CKB (Figure 3D and Table S1). This indicates that OPL2 may interact or form protein complexes with SOK3, AN, DLC1 and CKII and that these interactions can take place in the stomatal lineage (Figure 3E).

## Protein domain prediction software suggest that OPL2 interacts with SOK3 and DLC1 through Zn finger and beta-sheet motifs, respectively

Having identified DLC, SOK3, AN and CKII-subunit peptides enriched in all OPL2 samples, we sought to determine whether these proteins could directly interact with OPL2, and the nature of their protein-protein interfaces. OPL2 lacks clear functional motifs (see UniProt ID F4IRZ1), and overall, OPL2 secondary structure predictions with AlphaFold returned per-residue model confidence scores (pLDDTs) below 50, indicative of unstructured regions [41]. An exception was a finger-like fold comprised of cysteines C38, C50, C53 and histidine H41 with pLDDT scores of 85.72, 91.12, 89.65 and 83.62, respectively (Figure 4A) [41, 42]. Another metric for evaluating structural predictions is the predicted aligned error (PAE) which reports confidence in the relative position of two residues within the predicted structure; here, low values indicate high confidence and high values (up to 30) indicate a high expected position error [41, 43, 44]. While again the overall OPL2 protein structure is poorly predicted (Figure 4B), the PAE was lowest for the finger-like fold and adjacent α-helix (orange square in Figure 4B).

**Figure 4:**
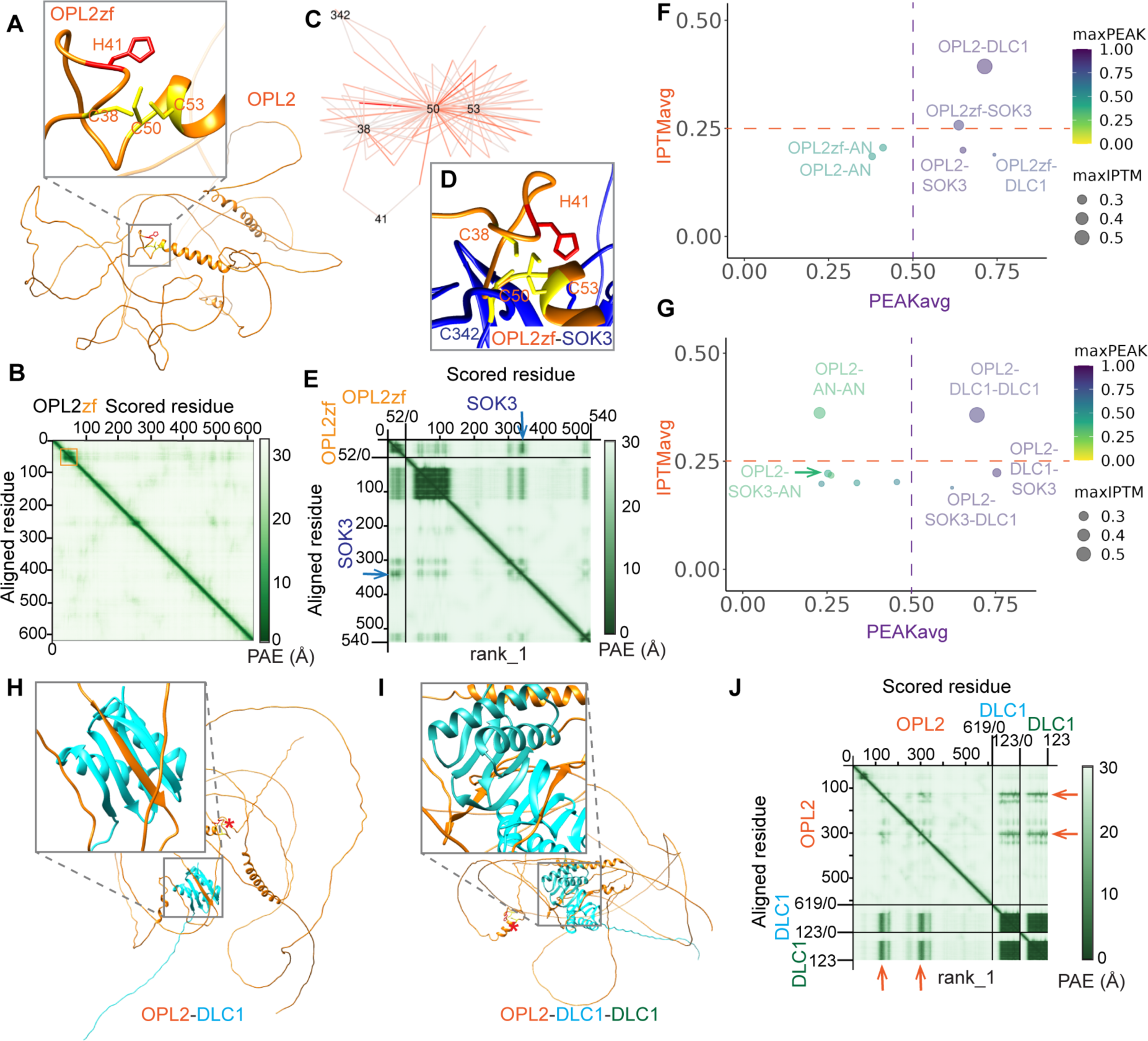
Protein structure predictions identify a putative Zn-finger in OPL2 and propose different interfaces for OPL2 interaction with SOK3 vs. OPL2 interaction with DLC1. (**A**) AlphaFold-predicted protein structure of OPL2 shows a largely disorganized conformation. (**B**) The Predicted Aligned Error (PAE) plot for OPL2 shows low confidence (light green) for the position of most OPL2 residues, except for a finger-like zinc finger motif (OPL2zf) marked by an orange square. (**C**) Protein Language Models and Attention Network Analysis (see methods) predicted a CHCC zinc finger motif in OPL2 comprised of residues C38, H41, C50 and C53. (**D**) AlphaFold-multimer prediction of OPL2zf (OPL2^P24-S75^ sequence of high confidence values in B) with SOK3 shows that the 3 zinc finger cysteines of OPL2^C38, C50, C53^ come in close proximity with SOK3^C342^, which together could create a zinc finger protein interface [45]. (**E**) Predicted Aligned Error (PAE) plot of OPL2zf with SOK3. Units: amino acid residues; light green to dark green: expected position error in Angstroms (**F-G**) AlphaFold-Multimer predicted ipTM scores (ipTM: interface predicted Template Modeling score) versus a custom PEAK score representing the inverted and scaled (0-1) minimum of the PAE between the protein chains, excluding intra-molecular interactions. For this study, predicted protein-protein interactions with an ipTM average >0.25, calculated across 5 different prediction models [46] and an average PEAK value >0.75 were considered high confidence and investigated in more detail. Dimers (F) using baits OPL2 and OPL2zf (OPL2^P24-S75^) versus preys DLC1, SOK3 and AN and trimers (G) using bait OPL2 versus all possible dimers of DLC1, SOK3 and AN were plotted (**H-J**) AlphaFold-multimer prediction of the OPL2-DLC1 interaction utilizes β-sheets of DLC1 to form a β-sheet complex with OPL2. DLC1 monomers (H) and dimers (I) interact with OPL2 via β-sheets, without involving the predicted OPL2 zinc finger (red asterisk). PAE plots represent OPL2 versus DLC1-DLC1 (J). Orange arrows point to high confidence score β-sheets of OPL2 that were induced the interface with DLC1 See also Figure S5

Intrigued by this potential finger-like OPL2 motif, we applied novel protein language models and attention network analysis (see methods), which revealed tight relationships between amino acid residues 38, 41, 50, 53 (and 342) (Figure 4C). Structural predictions and the interconnectedness of C38, C50, C53 and H41 support an OPL2 zinc finger (OPL2zf) motif comprised of three cysteines and a histidine (Figure 4A-C). Zinc fingers can mediate the formation of protein-protein interfaces, as cysteine residues bridge distinct proteins by forming a Zn^2+-^cysteine complex [45]. Using AlphaFold-Multimer models [41, 44, 46] we tested whether the OPL2zf domain (OPL2^P24-S75^) could scaffold interactions with our OPL2 proximity labelling candidates. With SOK3, we obtained a structural prediction that showed SOK3^C342^ in close proximity to OPL2’s C38, C50, C53 and H41 with corresponding PAEs of 7.68, 12.76, 5.24 and 7.98, respectively (Figure 4D-E). Considering that most residues of OPL2 show very low confidence with PAE values close to 30 (Figure 4B), these low PAE values of the OPL2-SOK3 zinc finger interface suggest a strikingly accurate prediction and suggest that OPL2 and SOK3 could interact via a Zn^2+^-complex. AlphaFold-multimer models running OPL2^P24-S75^ with other candidates AN, DLC1 and CKII, however, did not result in strong support for interactions mediated through the proposed OPL2zf. It is tempting to speculate that OPL2 protein interaction partners may each interact via distinct OPL2 domains and that the proposed OPL2zf is but one of many potential domains for protein interfaces.

To reveal other potential protein interfaces of OPL2 that may become only apparent during a specific protein-protein interaction, we ran AlphaFold-multimer predictions of full-length OPL2 with our top interaction candidates (Figure 4F). Interface pTM scores are a confidence metric that depicts the accuracy of protein-protein interfaces by scoring the interactions between residues of distinct protein chains [43, 44]. Our custom PEAK score uses the inverted and scaled (0-1) minimum of the PAE between the protein chains, excluding intra-molecular interactions. By plotting the average ipTM versus the PEAK scores of five distinct prediction models [46], we obtained confidence estimates for the interfaces between OPL2 as bait and AN, DLC1 or SOK3 as monomeric or dimeric preys (Figure 4F-G). Higher ipTM and PEAK scores depict better confidence in the protein interface. To separate our predictions into groups of lower and greater confidence, we chose a cut-off of ipTM >0.25 and PEAK >0.5 (Figure 4F-G, dashed lines). Of note, the ipTMavg score of 0.189 for OPL2zf-DLC1 was two times lower than the one of OPL2-DLC1 (ipTMavg = 0.393), which suggests that the hypothetical OPL2zf is not involved in binding to DLC1. Consistent with this hypothesis, the 3D structural prediction of the OPL2-DLC1 interface defines a β-sheet within the OPL2 structure that is absent in predictions of monomeric OPL2 (Figure 4H). This newly discovered OPL2 β-sheet is in close proximity to the highly conserved antiparallel β-sheets of DLC1 (Figure 4H). β-sheets interactions are the typical mode by which LC8-type DLCs serve as adaptors for scaffolding protein complexes [47, 48]. As expected, the protein interface of OPL2 with DLC1 dimers scored well and resulted in formation of two OPL2 β-sheets, each close to the β-sheets of one DLC1 but separate from the hypothetical OPL2zf (Figure 4I, OPL2zf is marked by a red asterisk). The PAE scores were particularly low for these OPL2 β-sheets (Figure 4J, orange arrows), which indicates high positional confidence within the protein structure.

Surprisingly, OPL2zf-SOK3 scored better than OPL2-SOK3 (Figure 4F) or any trimer of OPL2 with SOK3 (Figure 4G), which could argue for a transient Zn2+-interface between OPL2 and SOK3 that is only possible if OPL2 assumes a certain conformation. Although SOKs interact with AN [37], protein interface scores between OPL2-AN or OPL2-AN-SOK3/OPL2-SOK3-AN were poor (Figure 4F-G), suggesting that OPL2 does not directly interact with AN. We surmise that OPL2 may form potential protein-protein interfaces with DLC1 via β-sheets and with SOK3 via a putative zinc finger. Based on the confidence scores, a model where OPL2 interacts with either DLC1 or SOK3 has greater support than one where all three interact in a common complex.

We identified nearly all the catalytic (α, CKA) and regulatory (β, CKB) subunits of the tetrameric (2α2β) serine/threonine kinase CKII [21] among the peptides significantly enriched in OPL2 samples (Figure 3). OPLs are highly phosphorylated proteins [17], and OPS and OPL2 can be made hyperactive/stable by mutation of a conserved phosphosite [11, 17] suggesting that CKII could regulate OPL2 activity via phosphorylation. We used AlphaFold-multimer prediction models to determine which catalytic or regulatory subunits may form an interface with OPL2 (Figure S5), initially by predicting dimers between OPL2 and all of the 8 possible Arabidopsis CKII subunits. Catalytic subunit CKA3 had the highest ipTMavg score (0.427). CKB3, the weakest scoring protein, was the only CKII subunit not enriched in our OPL2 proximity labeling samples (Figure S5A and Table S1). We then modelled which regulatory subunits formed an interface with OPL2-CKA3 with highest confidence (Figure S5B). The CKII tetramer shows an extraordinary heterogeneity in plants and can utilize almost all combinations of CKA and CKB subunits [49]. We tested which regulatory subunits had the best predicted confidence to form an interface with OPL2-CKA3. Among all possible combinations, trimer OPL2-CKB4-CKA3 had the highest ipTMavg score of 0.386 and OPL2-CKA1-CKB1 the best PEAK score (Figure S5B). Protein structure models show predicted interactions between OPL2 and the CKA3 monomer (Figure S5C), or the dimers CKB4-CKA3 (Figure S5D) and CKB1-CKA1 (Figure S5E). However, PAE values between OPL2 and CKII subunits showed little confidence in strong protein interfaces (Figure S5F-H). We therefore conclude that a functional CKII tetramer utilizing any combination of subunits CKA1, CKA3, CKB4 and CKB1 is most likely to interact and potentially phosphorylate OPL2 to regulate its activity.

## Experimental validation of OPL interfaces with DLC1 and SOK3 indicates cell-cycle phase dependent interactions

Because the high degree of CKII pleiotropy and redundancy and between subunits makes genetic and protein co-localization studies extremely difficult [50, 51], we focused on the relationships between OPL2, DLC1 and SOK3. We tested the co-localization of OPL2 with DLC1 and SOK3 *in planta* using transient expression in *Nicotiana benthamiana* leaf epidermal cells. We generated translational fusion lines At*UBQ10p:*At*OPL2-mCherry,* At*UBQ10p:*At*DLC1-mGFP* and At*UBQ10p:*At*SOK3-mGFP*. Because OPL2 polarizes during interphase, but also shows dynamic polarity during cell division (Figure 1 and 2), we induced *N. benthamiana* epidermal cell divisions by co-infiltrating the leaves with At*CYCD3;1* [52]. When At*UBQ10p:*At*OPL2-mCherry* and phragmoplast microtubule marker *AtMAP65-4p:AtMAP65-4-GFP* [52] were expressed in this system and imaged between 48-72h post-infiltration (Figure 5A-D), non-dividing cells showed exclusively plasma membrane associated OPL2-mCherry signal, and the signal was polarized in multiple pavement cell lobes (Figure 5A). If cells were in cytokinesis, however, their phragmoplasts were marked by MAP65-4-GFP and OPL2-mCherry localized to the newly forming cell plate (Figure 5B-B’’ and 5D). As division completed, OPL2-mCherry signals were particularly strong at the plasma membrane adjacent to the new cell walls (Figure 5C-D), similar to OPL2 polarity in young GCs (Figure 1G).

**Figure 5:**
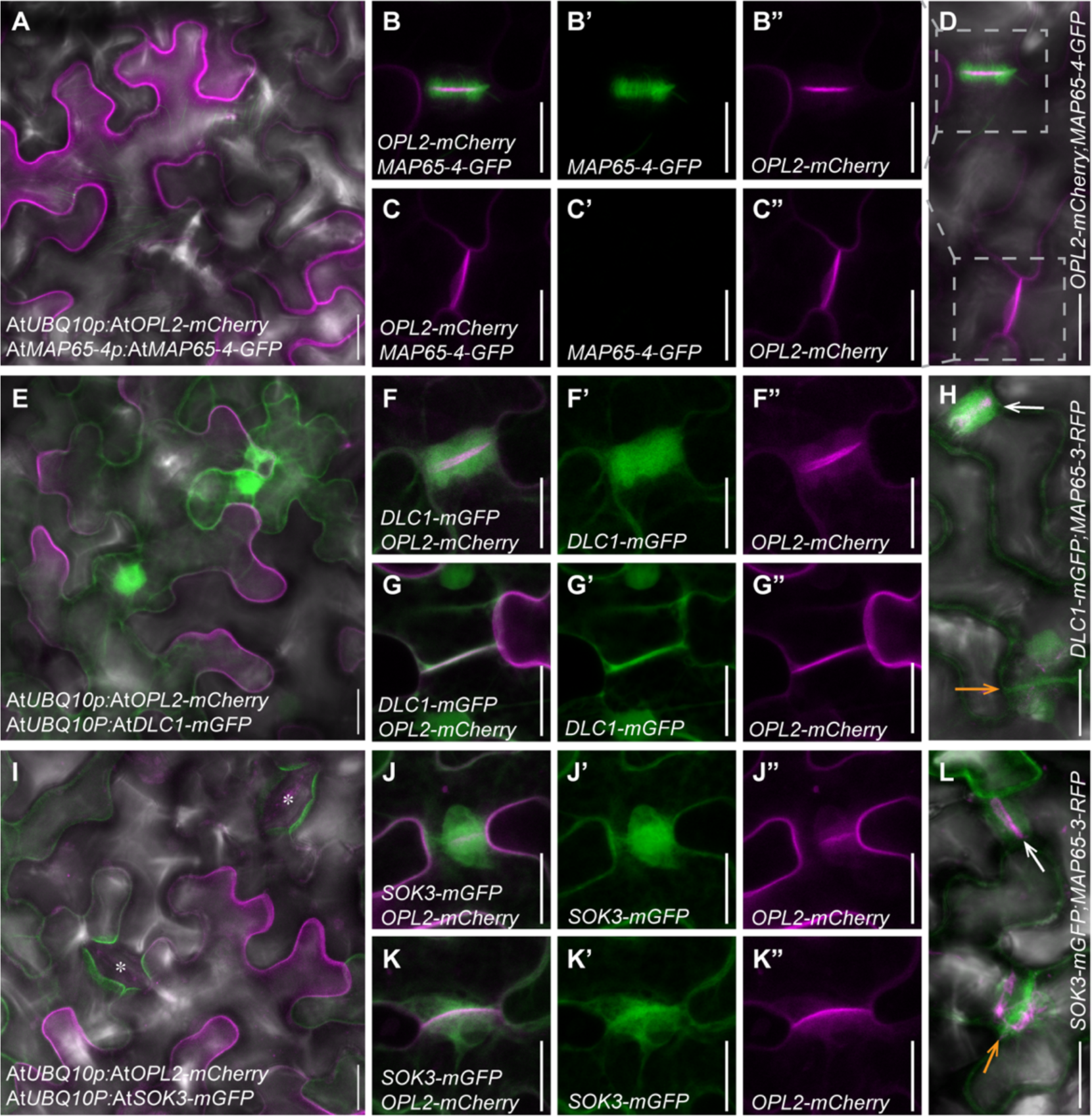
OPL2 co-localizes with microtubules, SOK3 and DLC1 at phragmoplast. (**A-D**) Transient expression of At*UBQ10p:*At*OPL2-mCherry* (magenta), phragmoplast microtubule marker At*MAP65-4p:*At*MAP65-4-GFP* (green) in *Nicotiana benthamiana* alongside At*CYCD3;1* to induce cell divisions [52]. OPL2-mCherry polarizes in lobes and dents of *N. benthamiana* pavement cells (A) and localizes at the forming cell plate during phragmoplast formation (merged in B, GFP channel in B’ and mCherry channel in B”) and at young, recently formed cell walls (C-C”). B and C are magnification of cell divisions in D (**E-G**) Transient co-expression of At*UBQ10p:*At*OPL2-mCherry* (magenta) and At*UBQ10p:*At*DLC1-mGFP* (green): PM polarized OPL2-mCherry localization and cytoplasmic/nuclear localized DLC1-mGFP do not overlap (E) unless cell divisions are induced by At*CYCD3;1* (F-G). OPL2-mCherry localizes to the growing cell plate, while DLC1-mGFP resides in the cell plate assembly matrix (CPAM) during the phragmoplast stage (merged in G, GFP channel in G’ and mCherry channel in G”). At fully formed but young cell walls, OPL2-mCherry and DLC1-mGFP co-localize at the PMs of the new walls (G-G”). (**H**) Co-expression of At*UBQ10p:*At*DLC1-mGFP* (green) with phragmoplast microtubule marker At*MAP65-4p:*At*MAP65-3-RFP* (magenta) also highlights the difference between DLC1-mGFP localization in the CPAM of phragmoplasts (white arrow) versus at the newly formed cell walls post cell division (orange arrow). (**I-K**) Transient co-expression of At*UBQ10p:*At*OPL2-mCherry* (magenta) and At*UBQ10p:*At*SOK3-mGFP* (green): PM polarized OPL2-mCherry localization and SOK3-mGFP polarity around mature GCs do not overlap (I) unless cell divisions are induced by At*CYCD3;1* (J-K). OPL2-mCherry localizes to the growing cell plate, while SOK3-mGFP resides in the cell plate assembly matrix (CPAM) during the phragmoplast stage (merged in J, GFP channel in J’ and mCherry channel in J”). At fully formed but young cell walls, OPL2-mCherry and SOK3-mGFP co-localize at the PMs of the new walls (K-K”). (**L**) At*UBQ10p:*At*SOK3-mGFP* (green) with phragmoplast microtubule marker At*MAP65-4p:*At*MAP65-3-RFP* (magenta) shows SOK3-mGFP localization in the CPAM of phragmoplasts (white arrow) versus at the newly formed cell walls post cell division (orange arrow). Scale bars 20 µm.

We next co-expressed At*UBQ10p:*At*OPL2-mCherry* and At*UBQ10p:*At*DLC1-mGFP*. In non-dividing cells, DLC1-mGFP was cytoplasmic while OPL2-mCherry polarized at the plasma membrane of pavement cells (Figure 5E). These proteins did not co-localize unless cell divisions were induced by At*CYCD3;1* [52]. During cytokinesis, OPL2-mCherry localized to the growing cell plate while DLC1-mGFP resided in the cell plate assembly matrix (CPAM) of the phragmoplast (Figure 5F-F’’). Within the CPAM, cell plate-forming vesicles are tethered and directed by molecular complexes before they fuse with the cell plate to maintain its growth [53]. As soon as cell division was complete, DLC1-mGFP and OPL2-mCherry both co-localized at the PM of the new cell walls (Figure 5G-G’’). To confirm that the “cloud” of DLC1-mGFP is truly associated with phragmoplasts, we co-expressed At*UBQ10p:*At*DLC1-mGFP* with phragmoplast microtubule marker *AtMAP65-3p:AtMAP65-3-RFP* [52] and indeed observed DLC1-mGFP first in the CPAM of phragmoplasts and later at the newly formed cell walls as division was completed (Figure 5H).

Finally, we co-expressed At*UBQ10p:*At*OPL2-mCherry* and At*UBQ10p:*At*SOK3-mGFP*. In non-dividing cells, OPL2-mCherry and SOK3-mGFP were plasma membrane associated and polarized, but they did not polarize to the same lobes (Figure 5I). When cell divisions were induced, however, SOK3-mGFP resided in the CPAM during phragmoplast formation (Figure 5J-J’’) and to the PM of newly formed cell walls post division (Figure 5K-K’’). Again, co-expression of SOK3-mGFP with phragmoplast marker MAP65-3-RFP confirmed these localization patterns during cytokinesis (Figure 5L). Interestingly, some OPL2-mCherry signal extended into the CPAM whenever co-expressed with DLC1-mGFP and SOK3-mGFP (Figure 5F’’ and 5J”), but not when expressed with MAP65-4-mGFP (Figure 5B’’). This suggests that SOK3-mGFP and DLC1-mGFP can recruit OPL2-mCherry to the CPAM. Altogether, the evidence supports OPL2 forming a complex with DLC1 and SOK3 during cell plate assembly.

### OPL2 and DLC1 co-localize at the leading zone of phragmoplasts during Arabidopsis stomatal lineage cell divisions

Polarized SOKs have been linked to the orientation of cell division planes [19, 37], but roles for plant DLCs in cell divisions, or in any cellular processes, are largely unknown. Dyneins are motors that transport cargo toward the minus end of microtubules and consist of heavy, intermediate and light chains plus adapter proteins [54]; however, only DLC proteins are present in Arabidopsis, and no cell polarity or division functions have been ascribed to them [20, 55]. Functional network analysis did suggest that DLC1 (At4g15930) could interact with microtubule associated proteins [20], including microtubule bundling factor MAP65-4 [56], and we confirmed co-localization of DLC1 with MAP65-4 in the phragmoplast (Figure 5H).

To obtain higher resolution images at native expression levels in the stomatal lineage, we generated *DLC1p:DLC1-YFP* and co-expressed this reporter with *TMMp:OPL2-BFP*. During both asymmetric and symmetric stomatal lineage divisions, DLC1-VENUS localized to the leading zone of phragmoplasts while OPL2-BFP occupied the membranes of the growing cell plate (Figure 6A-C). This was true for all GMC divisions (Figure 6A-A’’) as well as for asymmetric divisions of meristemoids (Figure 6B-B’’) and protodermal cells (Figure 6C-C’’), and became especially apparent in 3D reconstructions that showed a ring of DLC1-YFP along the growing edges of the cell plate (Figure 6A’-C”). Like the localization of DLC1 in the CPAM in *N. benthamiana*, we could again see a “cloud” of DLC1-VENUS in the CPAM of stomatal lineage phragmoplasts in z-projections (Figure 6D). We therefore conclude that OPL2 and DLC1 can only interact during cell divisions in the CPAM or the leading zone of phragmoplasts, where vesicles are fused to the growing cell plate. Given the predicted role of SOKs in orienting cell plates, we hypothesize that OPL2 forms a complex with DLC1 and SOKs to scaffold effectors but also bundle microtubules and/or regulate transport of cargo to the new cell walls during cell division (Figure 6E).

**Figure 6:**
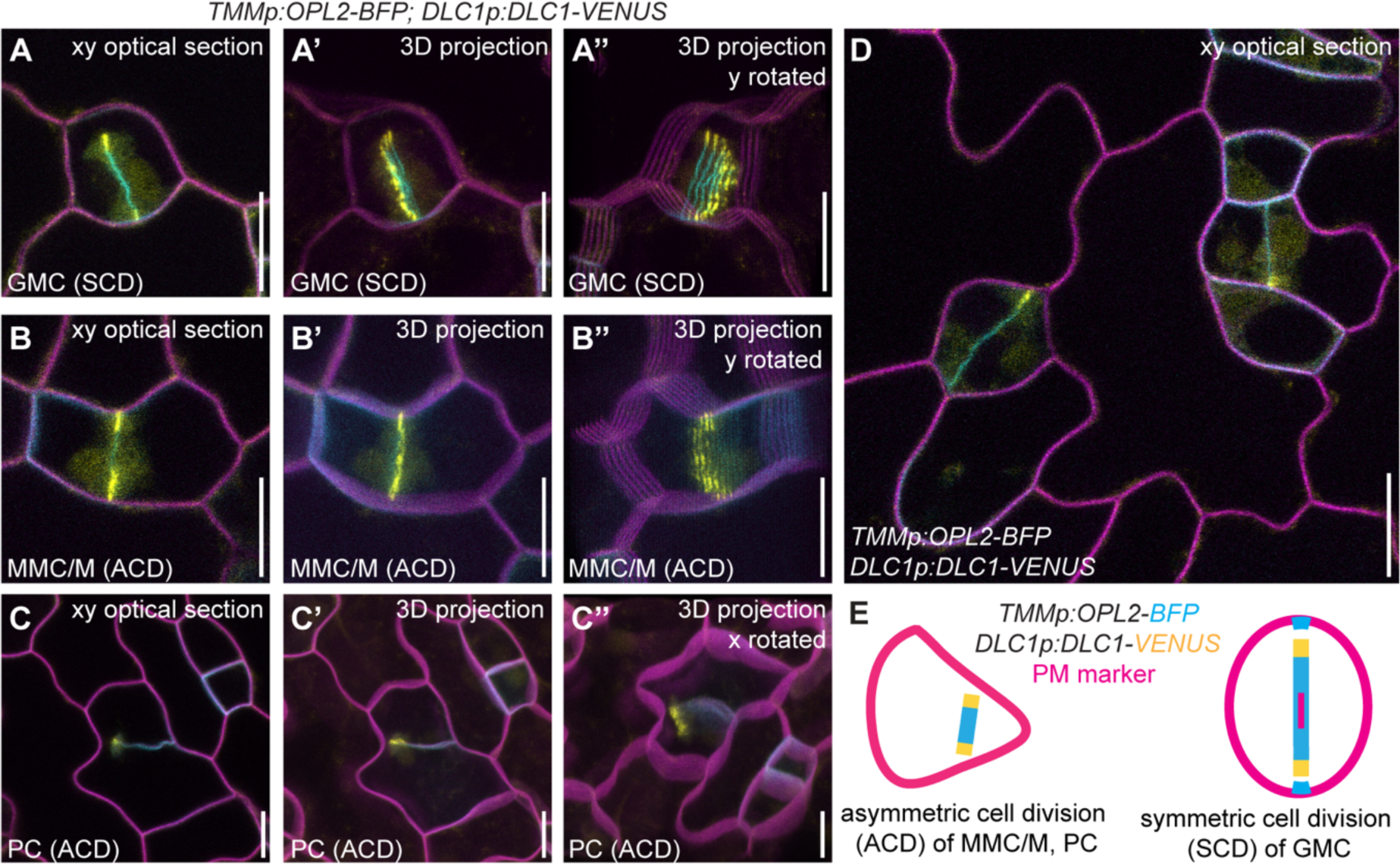
OPL2 co-localizes with DLC1 at the leading zone of phragmoplasts. (**A-D**) A triple marker line expressing *TMMp:OPL2-BFP* (blue), *DLC1p:DLC1-YFP* (yellow) and PM marker *ML1p:mCherry-RCI2A* is shown during symmetric cell division (SCD) of GMCs (A and D), asymmetric cell divisions (ACDs) of MMC or meristemoids (B and D) and ACD of a pavement cell (C). Depicted are optical sections through the cell in xy (A-C), 3D z-projections (A’-C’) and 3D projections rotated in y approximately 60° counter-clockwise to show the expanding cell plate (A’’-C’’). Scale bars represent 10 µm. (**E**) Schematic representation summarizes DLC1-YFP localization to the leading zone of the expanding phragmoplast in ACD and SCD with OPL2-BFP at the whole expanding cell plate and in polar patches marking the long axis during SCD.

## Discussion

The stomatal lineage presents some of the most fascinating uses of cell polarity in defining and patterning cell types and contributing to functionally important cell morphologies. The sparse and transient nature of stomatal lineage cells and their dependence on intrinsically disorganized scaffolding proteins also present some of the greatest challenges for *in vivo* proteomics. Here we demonstrate that proximity-labeling based proteomics and AI-guided protein structure predictions are powerful tools that can be adapted to reveal novel protein complexes and protein interfaces during cell polarity processes. In this work we also highlight the role of polarity proteins in the symmetric division that creates guard cells, showing how OPL2 localization predicts the division plane, marks regions that are microtubule enriched, and ensures the timely initiation of the stomatal pore.

The number of cellular compartments and processes to which biotin ligase-based proximity labeling has been successfully applied in plants continues to grow. This is, to our knowledge, the first application of the methodology to identify local proteomes of plasma-membrane associated proteins that reside in restricted polar subdomains. Compared to labeling nuclear proteomes of stomatal lineage cell types [27, 57], proximity labeling of polar domains was technically more challenging. We found that fusion to biotin ligases often disrupted function of polarity proteins and their mobility at the plasma membrane and when shuttling between the cytoplasm and plasma membrane [33, 58, 59]. We chose mini-TurboID (mTb) to minimize tag-induced disruption and tested all constructs in the stomatal lineage of stably transformed Arabidopsis cotyledons for their ability to complement mutant phenotypes and for their expected dynamic polar localization (Figure S3). We also used the shortest biotin labeling time (2h) possible to obtain signal but minimize unspecific cytoplasmic background labeling that increased over time. Consequently, the number of significantly enriched proteins we obtained was relatively low compared to other proximity labeling studies, and we did not recover all expected interaction partners of BRXL2 [5, 6].

We generated several transgenic lines that utilized distinct promoters (*OPL2p, BRXL2p, TMMp*) of different activity and tissue specificity. This enabled us to sample the full range of cell types in which OPL2 marks a polar domain, including proteins in proximity to OPL2 and BRXL2 in root apices and developing phloem [12, 14], which will be valuable candidates for future studies in those cellular contexts. This strategy also ensured that we had high confidence in the proteins identified repeatedly across all experiments. After filtering all proteomes against the wild-type background, we made stringent choices in comparing sample groups, demanding that candidate OPL2 interactors be enriched when compared to the proteomes of oppositely polarized BRXL2, as well as to an unpolarized plasma membrane control (RCI2A).

Because DLC1, SOK3, AN and CKII emerged consistently as being enriched in the OPL2 domain, including from samples obtained with the stomatal lineage specific *TMMp*-driven OPL2-mTb line, we had high confidence in these proteins as potential OPL2 interactors in the stomatal lineage. Following our proximity labeling approach with structure prediction models of protein-protein interfaces between OPL2 and the identified candidates helped us formulate valuable hypotheses about the distinct protein-protein interaction interfaces proteins can form with OPL2, and co-localization studies in planta strengthened the connections among OPL2, DLC1 and SOK3 during cytokinesis.

Our experiments indicate that angiosperm-specific OPLs interact with broadly conserved plant proteins, such as SOKs [19, 37], CKII [21] and DLCs [20], which is in line with our previous finding that ectopically expressed OPLs can affect growth of early diverging bryophytes (Wallner *et al.*, 2023). These interactions also highlight potential mechanisms by which OPLs guide cell division potential and differentiation through interaction with the cytoskeleton and microtubule-interacting proteins.

While a connection has been made between orienting cell division planes and polarizing SOKs [19, 37], little is known about DLCs in plants. The name DLC (dynein light chain) immediately confers a link to the cytoskeleton, as dynein motor protein complexes are well known for transporting molecular cargo toward the minus end of microtubules [54]. Extant Angiosperms, however, do not encode the dynein heavy chains or any of the other components that make up a functional dynein motor protein, and have retained only DLCs [55, 60]. Current thinking about DLC functions is that these proteins may retain some residual function as microtubule-interacting proteins and/or that DLCs are more generally used as scaffolding or oligomerization units [20, 47]. In support of microtubule association, functional network analysis with Arabidopsis DLC1 linked it to the microtubule plus-end binding protein END BINDING 1 (EB1) [61], phragmoplast microtubule bundler MAP65-4, [56] and Rho-GTPase family proteins implicated in many microtubule-dependent processes [62]. Consistent with a microtubule associated role, we showed that DLC1-VENUS localizes to the MT-rich leading zone of phragmoplasts during cytokinesis (Figure 5 and Figure 6). The other proposed role of DLCs became evident through protein structure prediction of the OPL2-DLC1 interface (Figure 4). LC8-type DLCs have the capacity to interact with an astonishing number of proteins that are not related to the dynein motor complex [63]. As adaptable linkers, LC8-type DLCs possesses a core of antiparallel β-strands that are highly ordered and conserved in all eukaryotes [48]. Their ordered β-strands form protein-protein interfaces by recognizing an 8-amino acid motif within intrinsically disordered regions (IDRs) of binding partners and induce structural re-organization of the IDR into an interacting β-strand [47, 63]. Consequently, LC8-type DLCs mediate structural organization and oligomerization of initially unstructured scaffolding proteins. Strikingly, DLC1-dependent induction of β-strands in OPL2 is exactly what we saw in our AlphaFold-multimer prediction models (Figure 4H-J). Likewise, SOK proteins have been proposed to mediate oligomerization of scaffolds and effectors as prerequisite for forming their polar domains [37]. It is therefore compelling to speculate that angiosperm specific OPLs can utilize highly conserved scaffolding and polarity-generating mechanisms to regulate cell divisions in the stomatal lineage [11].

Interestingly, SOK3 was predicted to have a high affinity for one of the few structured and confidently predicted regions in the OPL2 protein –the predicted zinc finger (Figure 4A-E). As zinc fingers serve multiple roles, including formation of protein-protein interfaces through binding of Zn^2+^ [45], this finding provides further evidence that OPL2 could acts in a polar scaffolding complex with several partners and effectors. While prediction models are powerful tools, they cannot yet capture all necessary parameters and signaling components present in a living cell. These missing parameters could induce conformational changes and thereby allow interactions between domains that cannot be predicted *in silico*. One example is the failure to find the experimentally confirmed interaction between SOK3 and AN [37]. We therefore do not want to rule out OPL2 participating in a larger complex with SOK3, AN and DLC1 *in vivo*. Another scenario may be that OPL2 oligomerizes with DLC1 and only binds to or recruits SOK3 and its effector AN during stomatal lineage cell divisions. While our co-localization studies can restrict OPL2-DLC1-SOK3 multimer formation to the CPAM and leading zone of phragmoplasts (Figure 5 and Figure 6), further studies are needed to decipher the possible combinations of proteins in this complex, including its spatio-temporal assembly and role in plant cell divisions.

Our data could also shed light on the mysterious role of SOKs in conferring polarity and orienting cell divisions [19, 37]. The evidence that SOKs could be polar scaffolding complexes is compelling, but thus far only AN, a protein of highly debated function, has been identified as a SOK partner [19, 37]. Moreover, the DIX domain of SOKs is proposed to be the central hub for plant cell polarity complexes and famously known for being the only DIX domain in plant proteins [37]. In animals, Dishevelled (Dsh) possesses a DIX domain to confer planar polarity through polymerization as integral component of Wnt signaling [64]. We see parallels between SOK/OPL dynamics in plants and those of Dsh/Prickle (Prk) in animals. In Drosophila epithelial cells, Prk regulates planar cell polarity by antagonizing Dsh through binding of the Dsh DEP domain, which prevents Dsh cortical localization [65]. Integral to Prk and its role in regulating cortical polarity are the LIM domains that contain two zinc fingers each [66]. Although OPL2 does not possess LIM domains, the predicted zinc finger could serve a similar function in mediating protein-protein and cytoskeleton interactions [67]. Likewise, CKII, a protein kinase conserved in all eukaryotes [21], can phosphorylate Dsh [38] but is also itself repressed by Dsh2 and activated by Prk [68] (Figure 7).

**Figure 7:**
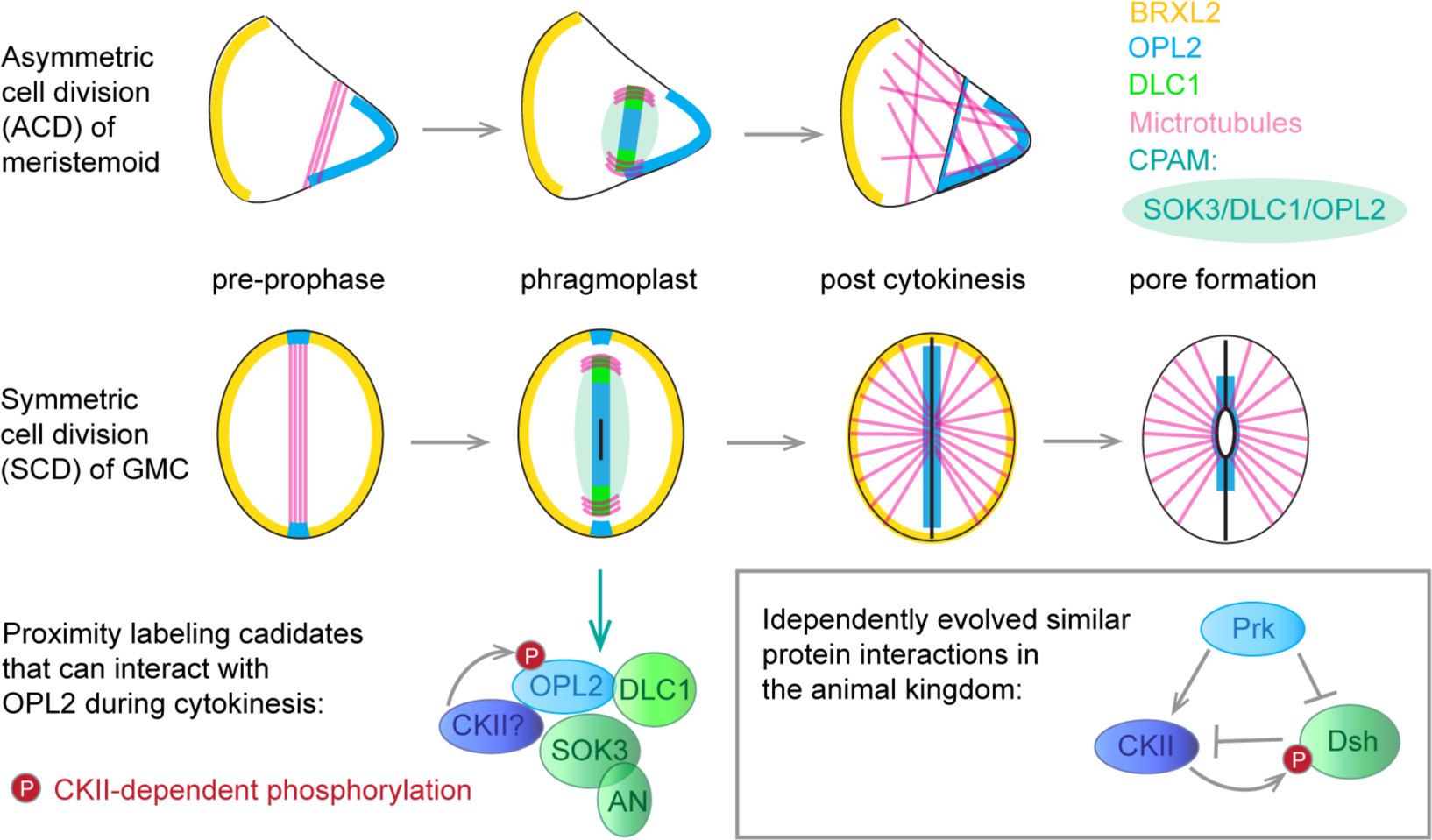
Model for OPL2 and partners during cell division and stomatal pore formation. The subcellular distribution of polarity proteins and partners during critical stomatal lineage divisions is cartooned, with a special emphasis on the relationships to microtubule (pink) distribution at different cell cycle phases and during stomatal differentiation. The proposed complex of polarity proteins as defined by proximity labeling and structural modeling is at the bottom, and a comparison to animal protein complexes involved in division orientation and cell polarity boxed [38, 65, 67, 68]. In both systems, IDR-containing molecules assemble into scaffolding complexes that have the potential to interact with intracellular signaling systems.

CKII has many targets throughout eukaryotes and was enriched in our OPL2 proximity-labeling samples (Figure S3). Although catalytic α-subunits CKA are typically nuclear localized or chloroplast-specific, both CKB1 and CKB4 are reported to localize to the cytoplasm in plants [49], which could allow targeting of or by OPL2. CKA1 and CKB1, our top hits after employing AlphaFold-multimer prediction models (Figure S5), form an especially active CKII in Arabidopsis [69]. Consequently, CKII could phosphorylate OPL2 (as we argue below), but also act downstream of OPL2 or be regulated by OPL2, similar to the situation with CKII and Prk [68]. Although the plant and animal kingdom evolved separately, it is fascinating to compare how proteins from different kingdoms with structurally similar domains behave. If similar principles are employed between animal and plant cortical polarity, we may well find similar protein domains and principles in establishing cell and tissue polarity.

OPLs are highly phosphorylated proteins, and their activity is modulated by phosphorylation at a conserved Q-**S**-E-I-A-D motif [11, 17], which is similar to the CKII recognition motif S/T-D/E-x-D/E [70]. It is therefore likely that CKII can phosphorylate OPL family members (Figure S5). We showed, using a OPL2^S295K;E296K^ (OPL2m) phosphosite variant considered to render OPL2 hyperactive/stable [17], that prolonged OPL2 persistence at the ventral wall of developing guard cell complexes suppresses stomatal pore formation (Figure 2 and Figure S2). Although we can only speculate about reason for this pore-suppression, it is quite clear that OPLs mark polar domains that coincide with preprophase band formation prior to GMC division and are strongly attracted to the new ventral walls of the young GCs (Figure 1 and Figure 2).

We recently reported that the BRXL2-BASL domain excludes microtubules to guide nuclear migration and division plane positioning during asymmetric stomatal divisions [10, 24]. OPL polarity domains, in contrast, are present in especially microtubule-rich regions (Figure 2 and Figure S2). Microtubules and specialized cell wall composition play roles in stomatal pore formation, and microtubules have been shown to regulate cellulose deposition by guiding cellulose synthases to the right cell wall domains [71, 72]. We could therefore speculate that OPL2 localizes to the growing cell wall to either delay pore formation until GCs have matured or arrange or interfere with microtubules and cellulose deposition [22, 72]. This question, including why OPLs and BRXLs occupy such distinct domains throughout stomatal development and scaffold very specific interaction partners (Figure 7), opens up exciting avenues for future studies of stomatal development and for the establishment of plant cell polarity in general.

### Limitations of the study

Genetic studies have been challenging for the BRXL, OPL, and SOK protein families due to redundancy although *BRX* quadruple and *OPL* quadruple mutants have been shown to have roles in the stomatal lineage [11, 19, 20, 37, 73]. The CKII and DLCI families feature even more family members, and transcriptional profiles suggest that they are broadly expressed. We generated CRISPR/Cas9 truncation alleles of *DLC1* but neither the single mutant, nor the mutant in combination with an *OPL2* null produced any clear phenotypes in stomatal development. In addition, while the advances in proteomics and structure prediction we used here accelerated understanding of potential roles of OPL2, it is still a highly disordered protein, and it has been recalcitrant to purification for its use in phosphorylation assays with CKII. Therefore, a complete picture of the role of OPL2 partners during cytokinesis inside and outside of the stomatal lineage will require additional studies.

## STAR METHODS

### Key resources table

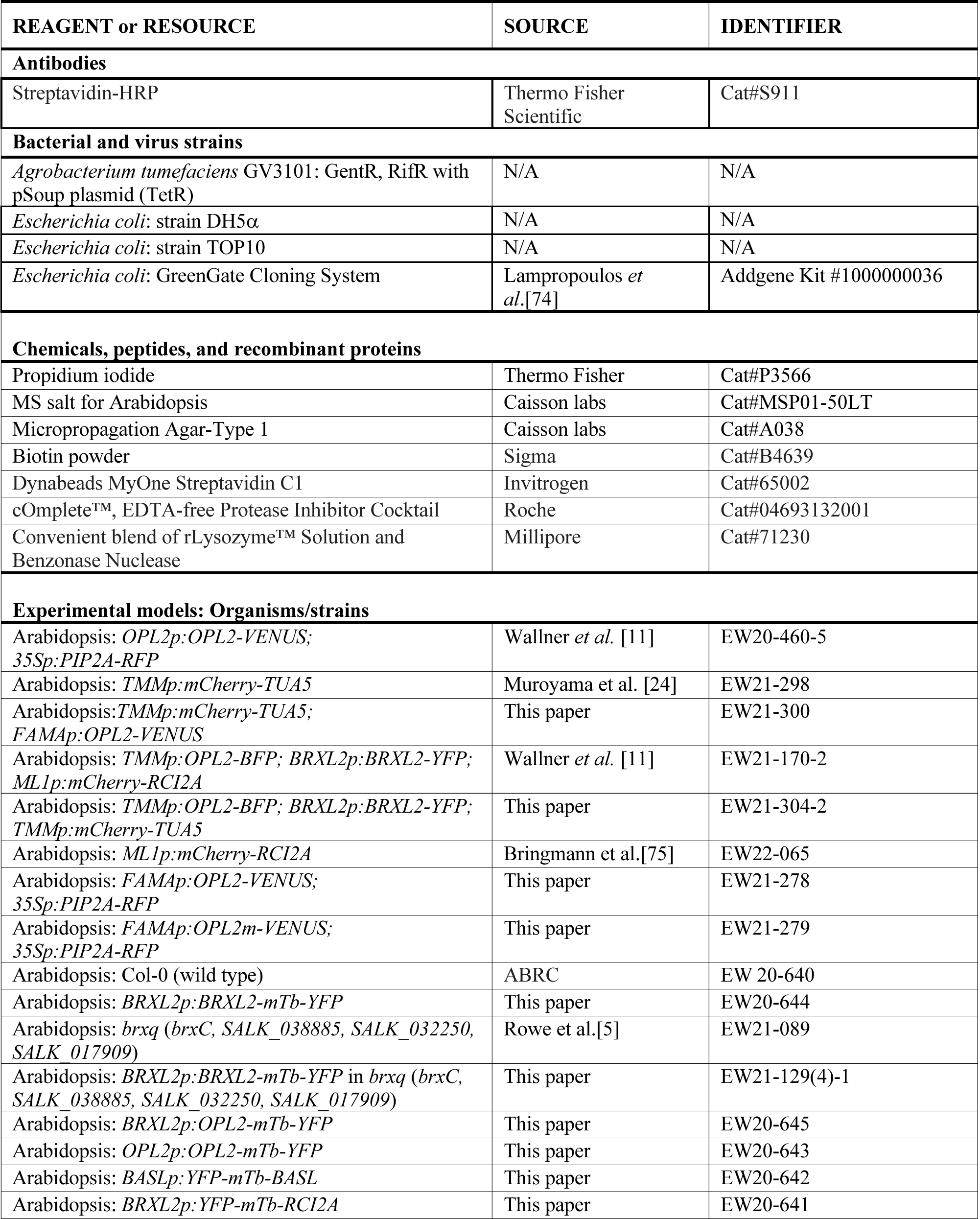
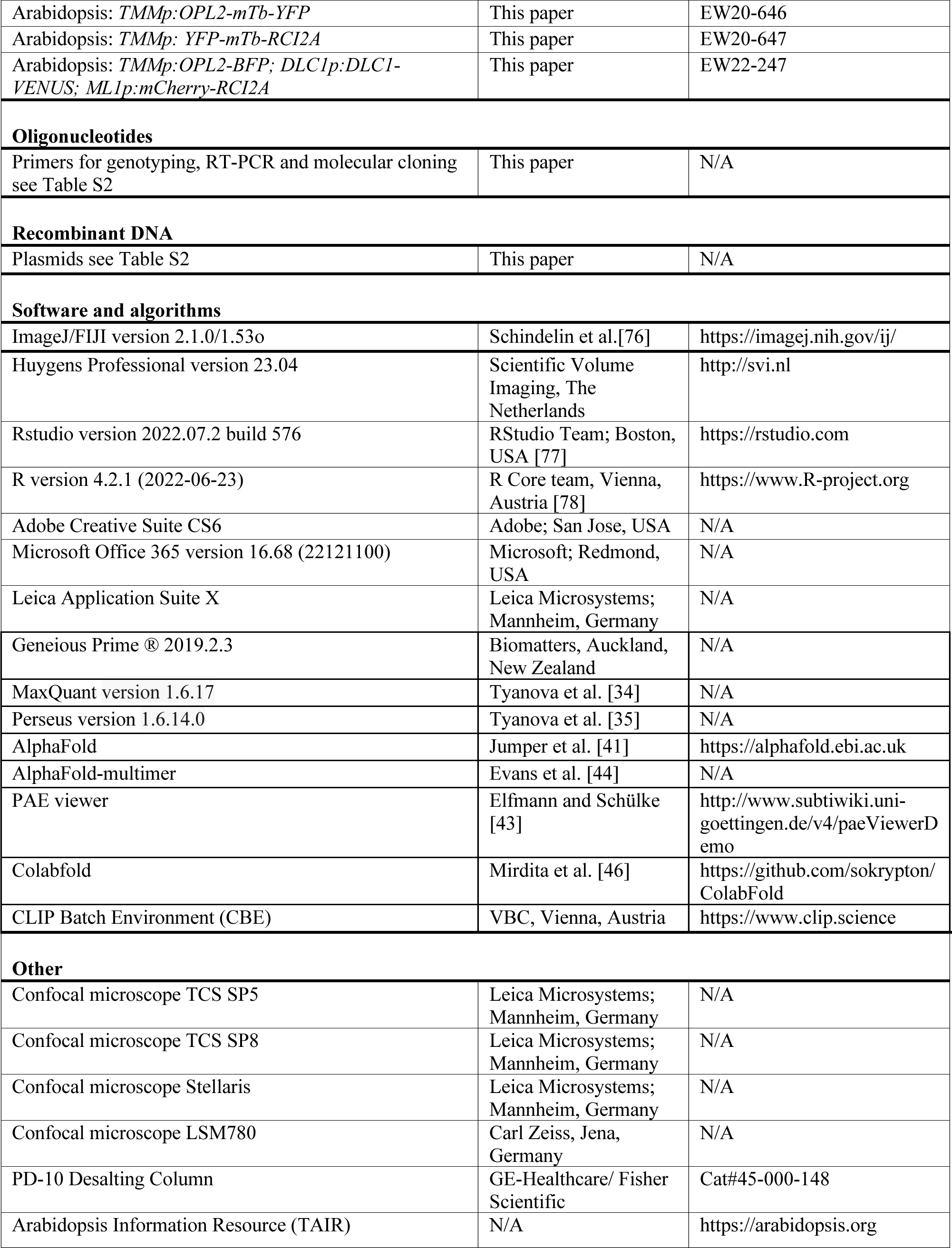

## RESOURCE AVAILABILITY

### Materials availability

Newly generated lines and sources of previously reported transgenic lines are listed in the Key resources table.

### Lead Contact

Further information and requests for resources and reagents should be directed to and will be fulfilled by the Lead Contact: Dominique Bergmann

### Data and code availability

All data required to substantiate the claims of this paper are included in main or supplemental data. Full proteomics dataset have been deposited in the PRIDE database and will be available immediately publication of this paper’s version of record. This paper does not report original code.

Any additional information required to reanalyze the data reported in this paper is available from the Lead Contact upon request.

## EXPERIMENTAL MODEL AND STUDY PARTICIPANT DETAILS

*Arabidopsis thaliana* ecotype Columbia (Col) was the genetic background for all mutants and transgenic lines. Specific genotypes are listed in the Key Resource Table. Seedlings were grown vertically in growth chambers with long day conditions (16 h light and 8 h dark) at either 23 °C day/21 °C night with blue, red and white LEDs at 109 µmol/m^2^s light intensity (GMI, Austria) or at 21 °C day/night with full-spectrum fluorescent tubes emitting 110 µmol/m^2^s light intensity (Stanford, USA) for 3-10 days. For each experiment, all plants were grown at the same time, location and under the same conditions alongside their own internal controls to ensure samples could be compared. All seeds were liquid sterilized in 70 % ethanol with 0.2 % Tween-20 for 5 min, washed twice with 100 % ethanol and air dried under sterile conditions. Sterile seeds were stratified in sterile water at 4 °C in the dark for 48 h and sown in rows directly on agar-solidified ½ strength Murashige and Skoog (MS) plates with 0.8 % plant agar and supplemented with 0.5 % sucrose for phenotyping and imaging.

## METHOD DETAILS

### Generation and origin of transgenic and mutant lines

Arabidopsis plants were transformed with plasmids listed in Table S2 by the floral dip method described in [79]. Transformed seedlings were selected on ½ MS plates with 0.8 % plant agar but without sucrose and the respective antibiotic/herbicide (7.5 mg/L sulfadiazine, 50 mg/L kanamycin, 50 mg/L hygromycin, 15 mg/L phosphinothricin) as listed in Table S2.

### Generation of plasmids for plant transformation

Translational fusion constructs were generated using the Green Gate method [74]. Gene sequences from genomic or cDNA were amplified by Phusion PCR with Green Gate-specific primers listed in Table S2 and cloned via BsaI restriction sites into entry modules with a pUC19 based vector backbone. Final constructs (Table S2) with pGreen-IIS based vectors backbone were generated by assembly of entry modules according to the Green Gate method [74].

### Imaging and tissue staining

The epidermis of abaxial cotyledons was imaged at 3, 4 or 7 days post germination (dpg) unless otherwise indicated in the figure legend. If not expressing a genetically encoded plasma membrane marker (Key Resource Table), cell walls were stained by incubating seedlings for 5 min in propidium iodide (PI, 40 µg/ml) dissolved in water supplemented with 0.02% of the detergent SILWET-L77 with subsequent rinsing in DI water for 2 min prior to imaging cell outlines. For visualizing PI staining and translational fusion proteins with fluorescent tag, a Leica SP5, SP8 or Stellaris confocal microscope with HyD detectors (Leica) or a LSM780 (Zeiss) were used with 25x or 40x water objectives. Lines expressing YFP (Yellow Fluorescent Protein) or VENUS were excited by an argon (SP5, SP8, LSM780) or white light laser (Stellaris) at 514 nm and the emission was detected between 520-540 nm. BFP (Blue Fluorescent Protein) was excited by UV at 405 nm, with emission collected between 420-470 nm. mCherry/PI and RFP were excited at 561 nm (DPSS) and emission was collected between 600-650 nm. All images displaying microtubules were deconvoluted using standard automatic settings in Huygens Professional (Scientific Volume Imaging).

### Proximity labeling

500 µl of seeds were harvested for the wild type control and for each transgenic plant line (*BRXL2p:BRXL2-mTb-YFP, BRXL2p:OPL2-mTb-YFP, OPL2p:OPL2-mTb-YFP, BASLp:YFP-mTb-BASL, BRXL2p:YFP-mTb-RCI2A, TMMp:OPL2-mTb-YFP, TMMp:YFP-mTb-RCI2A*, see Key Resource Table), surface sterilized and stratified in water (see plant growth conditions). Each batch of imbibed seeds was equally distributed on 6 ½ MS plates containing 0.8 % plant agar and 0.5% sucrose (ca. 100 µl seeds per plate) topped with filter paper (Whatman Shark Skin Filter Paper, GE Healthcare #10347509). Seedlings were grown on the filter paper-topped MS agar plates in a growth chamber under long-day conditions, 110 µmol/m^2^s light intensity for 4 days. 4-day-old seedlings were carefully taken off the filter paper. Seedlings from two plates of the same line were pooled into one technical replicate each and submerged in ½ MS liquid medium containing 50 µM biotin for 2 h in the long day growth chamber. After biotin treatment, the liquid was strained off and seedlings were rinsed with ice cold water (4°C) to stop biotinylation and washed 3x with 200 ml ice cold water for 2 min each to remove excess biotin from the plant tissue. Washed seedlings were patted dry, immediately frozen in liquid nitrogen and ground into a fine powder using a mortar and pestle.

### Western blotting

To test biotinylation efficiency, 100 µl of frozen plant powder was boiled in 200 µl 4x Leammli buffer (240 mM Tris pH 6.8, 8% SDS, 40% glycerol, 10% beta-mercaptoethanol, 0.01% bromphenol blue) at 95°C for 5 min. 5 µl of the supernatant was loaded onto a 10% SDS-PAGE and proteins were separated by 30mA until the bromphenol blue left the gel. Proteins were blotted onto Immobilon-P PVDF membrane (0.2 µm, Millipore) using a Trans-Blot Semi-Dry transfer Cell (BioRad) at 20 V for 90 min. Membranes were blocked in 5 % BSA in TBS-T for 1 0.2 µg/ml Streptavidin-HRP (S911, Thermo Fisher Scientific) in 5 % BSA in TBS-T at 4 °C overnight. Signals were detected on a ChemiDoc MP Imaging System (BioRad) with exposure time set to 30 sec.

### Protein extraction

3.5 ml of densely packed plant powder was aliquoted per sample (and technical replicate) into 15 ml falcon tubes and mixed with 2 ml protein extraction buffer (50mM Tris pH 7.5, 150mM NaCl, 0.1% SDS, 1% NP-40, 0.5% Na-deoxycholate, 1mM EDTA, 1mM EGTA, 1mM DTT, 20 µM MG-132, 1x cOmplete protease inhibitor mix (Roche), 1mM PMSF), vortexed for 30 sec and incubated for 15 min at 4°C on a rotor wheel. 1 µl Lysonase (Millipore) was added per sample to digest cell walls and samples were incubated for another 15 min at 4°C before sonicated in an ice bath 4x for 30 s on high setting using a Bioruptor UCD-200 (Diagenode) with 90 sec breaks on ice. Samples were centrifuged for 15 min at 4 °C, 15000 g and the supernatants were sent through PD-10 desalting columns (GE-Healthcare). Columns were equilibrated and samples were eluted with equilibration buffer (50mM Tris pH 7.5, 150mM NaCl, 0.1% SDS, 1% NP-40, 0.5% Na-deoxycholate). The protein concentration das measured of each eluate sample in a 1:20 dilution by Bradford (BioRad protein assay). Per sample, 15 mg of protein were transferred into a new 5 ml LoBind tubes (Eppendorf) and mixed with 200 µl Dynabeads MyOne Streptavidin C1 (Invirtogen) slurry was washed with extraction buffer on a magnetic rack and added to each eluate. 1xcOmpletee protease inhibitor mix and 1 mM PMSF were again added to each sample and extracts were incubated with Dynabeads for 16h at 4°C on the rotor wheel. Dynabeads were separated on a magnetic rack in 1.5 ml LoBind tubes (Eppendorf) and washed 4x with 1.5 ml cold equilibration buffer. On the last wash step, beads were transferred into a new LoBind tube and washed 1x with cold wash buffer I (equilibration buffer with 250 mM NaCl + 1% SDS), 1x with cold equilibration buffer, 2x with cold wash buffer II (equilibration buffer with 250 mM NaCl), 2x with cold wash buffer III (equilibration buffer with 500 mM NaCl), 1x with cold equilibration buffer and 2x with Urea wash buffer (2M Urea, 50 mM Tris pH 7.5, room temperature). Beads were resuspended in 1 ml cold 50 mM Tris pH 7.5 and transferred into a new 1.5ml LoBind tube and all buffer was removed using a magnetic rack. Beads were frozen at −80°C for further processing.

### Mass spectrometry sample preparation

Trypsination of peptides and mass spectrometry was exactly done as previously described in [27]. In brief, beads were washed 2x with 1ml 50mM Tris pH 7.5, 2x with 1ml Urea wash buffer III (50mM Tris pH 7.5, 1M Urea) and resuspended in 80µl Trypsin buffer (50mM Tris pH 7.5, 1M Urea, 1mM DTT, 0.4µg Trypsin). Samples were incubated for 3h at 25°C, rocking at 1200 rpm. The beads were washed 2x with 60µl Urea wash buffer III and the supernatant was combined with the trypsin digest supernatant. The eluate was first reduced by 4 mM DTT (25°C for 30min, 1000 rpm) and then alkylated with 10 mM iodacetamide (incubated at 25°C for 45min, 1000 rpm). 0.5µg trypsin were added to complete the digestion by incubating at 25°C over night (18 h, 800 rpm). Formic acid was added to a final concentration of 1 %. OMIX C18 desalting tips were activated by aspirating 2x 200µl buffer B2 (0.1% formic acid, 50% acetonitrile) and equilibrated by 4x 200 µl buffer A2 (0.1% formic acid). Peptides were bound to the C18 tips by aspirating and dispensing the peptide sample 8x, washed 10x with buffer A2 and eluted in 200µl buffer B2. Samples were dried in the speed vac and resuspended in 0.1% formic acid to be analyzed on a Q-Exactive HF hybrid quadrupole-Orbitrap mass spectrometer (Thermo Fisher), equipped with an Easy LC 1200 UPLC liquid chromatography system (Thermo Fisher), see [27].

### Data analysis with MaxQuant and Perseus

Protein identification and label-free quantification (LFQ) were done in MaxQuant (version 1.6.17) [34] using default settings, with data normalization done in MaxQuant through the LFQ algorithm [80]. A maximum of 5 modifications per peptide were allowed and trypsin/P with a maximum of two missed cleavages was set as digest. Peptides were searched against the latest TAIR10 protein database (TAIR10_pep_20101214) [81], including a custom list of contaminants, such as trypsin, human keratin, streptavidin, YFP, mTb-YFP alongside the list of contaminants provided by MaxQuant. Peptides with a minimum length of 7 amino acids and FTMS and TOF MS/MS tolerance of 20 ppm and 40 ppm, respectively and a peptide and protein FDR of 0.01, including unique and razor peptides. LFQ minimum ratio count was set to 1, with a minimum of 3 and average number of 6 neighbors. Raw data and MaxQuant and Perseus analysis is deposited to the ProteomeXchange Consortium (http://proteomecentral.proteomexchange.org) via the PRIDE partner repository [82] with the dataset identifier XXXXX. Data analysis with Perseus was done as described before [27]. In brief, the MaxQuant output file ‘proteinGroups.txt’ was imported into Perseus using the LFQ intensities as Main category. Proteins marked as ‘only identified by site’, ‘reverse’ and ‘potential contaminant’ (including trypsin) were removed from the data matrix. LFQ values were log2 transformed and were filtered based on valid values of minimum 2 in at least one group and missing values were replaced by the Perseus algorithm. First, all sample groups were compared to the wild type (as single control group) by a one-sided Welch’s t-test with FDR of 0.5 and S0 of 0.1 to filter out all unspecific background labeling. After all filtering steps, statistical comparisons between all other sample groups were displayed as Volcano plots, choosing the following parameters: Test = t-test, sides = both, no. of randomizations = 250, FDR = 0.5 and S0 = 0.1.

### AlphaFold and AlphaFold-multimer prediction pipeline

AlphaFold was used to predict the protein structure of OPL2 (AT2G38070, UniProt ID: Q8GW92) [41]. Alphafold-Multimer [41, 44, 46] was used to predict interactions between the ‘bait’ proteins and ‘candidate’ proteins with a script provided by D. Handler23 using mmseqs (git@95e1c10) for local MSA creation and colabfold(git@98a3d31) for structure prediction, which was run on the CLIP Batch Environment (CBE) cluster (see Key Resource Table). All protein sequences were obtained from The Arabidopsis Information Resource TAIR [81] for OPL2 (AT2G38070), OPL2zf (OPL2^P24-S75^), DLC1 (AT4G15930), SOK3 (AT2G28150), AN (AT1G01510), CKA1 (AT5G67380), CKA2 (AT3G50000), CKA3 (AT2G23080), CKA4 (AT2G23070), CKB1 (AT5G47080), CKB2 (AT4G17640), CKB3 (AT3G60250), CKB4 (AT2G44680) and fed into the pipeline in FASTA format. Predicted interaction confidence was determined using the Interface predicted TM-score (ipTM) and a custom PEAK score. The PEAK score was calculated as the inverted & scaled (0-1) minimum of the predicted aligned error (PAE) between the chains excluding intra-molecular interactions. PAE plots were analyzed using [43]. Five independent ipTM and PEAK scores were averaged, corresponding to the output of the five distinct prediction models AlphaFold2 (default settings with templates), AlphaFold-Colab (no templates), ColabFold-RoseTTAFold-BFD/MGnify, ColabFold-AlphaFold2-BFD/MGnify and ColabFold-AlphaFold2-ColabFoldDB [46].

### Identification of a Zinc-finger motif in OPL2 using Protein Language Models and Attention Network Analysis

To identify candidate motifs in OPL2, we designed a novel analysis leveraging ProtT5-XL-UniRef50 [83], a transformer-based protein language model pre-trained on a large corpus of protein sequence [84]. Protein language models contain hundreds of “attention” heads, or small networks that capture many types of relationships between amino acid in a sequence [85], including structural motifs. We reasoned that perturbing amino acids in the same motif would elicit similar patterns of disruption to specific attention networks. In a computational framework similar to Deep Mutational Scanning [86], we systematically introduced single-residue mutations across the entire OPL2 sequence. Following each mutation, we recorded a similarity score of each head’s attention network for the variant sequence compared to the original sequence.

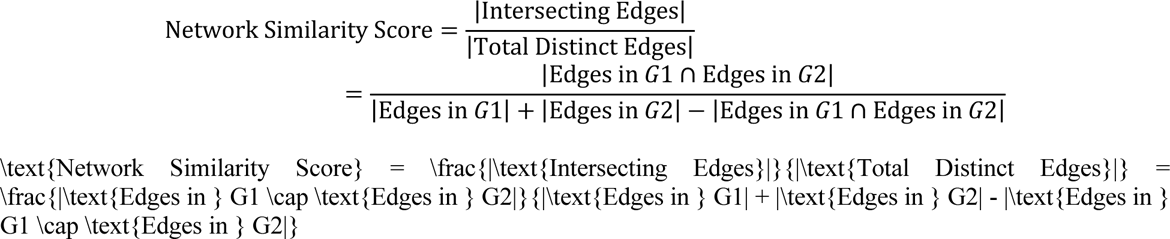

As the magnitude of similarity scores for most variants at the same position in the sequence was similar, we next averaged similarity scores for each position. These similarity scores were then transformed into Z-scores, benchmarked against the distribution of all similarity scores for the same head across the entire spectrum of mutated sequences. For our network visualization, we established connections (edges) between variant positions and specific attention heads whenever a variant’s Z-score for a given head fell below −6. Notably, amino acids C38, H41, C50, and C53 disrupted overlapping sets of attention heads (Figure 4C). Upon comparison to the OPL2 structure, these residues appeared to constitute a Zinc-finger motif. Code is available at https://github.com/clairemcwhite/attn_head_scan along with an interactive Colab notebook.

## QUANTIFICATION AND STATISTICAL ANALYSIS

Plasmid maps and sequence assemblies were performed in Geneious Prime (Biomatters). Gene sequences were obtained from The Arabidopsis Information Resource (TAIR) [81]. Confocal images were analyzed in ImageJ/FIJI [76]. Pore lengths and stomatal density measurements were statistically analyzed in R [78] and Rstudio [77] version 2022.07.2+576 and plots were created with the ggplot2 [87] package. Other packages used: tidyverse, ggbeeswarm, scales, ggpubr, knitr, ggsignif, FSA, ggstatsplot, rcompanion and multicompView (for detailed information see www.r-project.org). Datasets were first tested on normal distribution by a Shapiro-Wilk Test and homogeneity of variances by a Levene-Test (α=0.05). As data was not normal distributed, a Kruskal-Wallis multiple comparison followed by a post hoc Dunn test with p-values adjusted by the Holm method (p-value <0.05) was applied. Presented is one of 3 independent replicates. The type of statistical test used, the statistically distinct groups or p-values as well as the sample sizes (n) and their nature (e.g. individual cotyledons or cells observed) are indicated in the figure legends. The precision measures of all box plots are medians flanked by lower and upper hinges corresponding to the 25^th^ and 75^th^ percentile, respectively. Box plot whiskers extend from each hinge to the largest and smallest value measured with maximal 1.5-times the inter-quartile range. See further information on the geom_boxplot function from the ggplot2 package [87]. Precision measures of all other types of plots are described in the respective figure legends. Software is also referenced in the Key Resource Table.

## Supporting information

Proteomics data file

## Acknowledgements

We are grateful to Vienna BioCenter Core Facilities for access to plant growth chambers, microscopes and sequencing (VBC, Vienna, Austria). We would especially like to thank the BioOptics facility (IMP, Vienna, Austria) for training on microscopes and deconvolution software. Andi Pauli (IMBA) for access to the local AlphaFold-Multimer cluster. Natalie Edelbacher and Katharina Jandrasits for helping with plant potting and booking of growth chambers. We thank Bo Liu (UC Davis) for sharing plasmids for tobacco infiltration.

## Author contributions

ESW contributed conceptualization, data acquisition, analysis and interpretation, writing of the original draft and revision. ESW, AM, SX. contributed data acquisition and analysis. DH contributed software. LD contributed funding and manuscript revisions. DCB contributed supervision and funding, writing of the original draft and revision.

## Data and materials availability

All data needed to evaluate the conclusions in the paper are present in the paper and the Supplementary Materials. Proteomics data will be deposited in PRIDE with access granted immediately upon publication of this manuscript’s version of record.

## References

1. Goldstein, B., and Macara, I.G. (2007). The PAR proteins: fundamental players in animal cell polarization. Developmental cell 13, 609–622.

2. Wallner, E.S. (2020). The value of asymmetry: how polarity proteins determine plant growth and morphology. Journal of experimental botany 71, 5733–5739.

3. Kania, U., Fendrych, M., and Friml, J. (2014). Polar delivery in plants; commonalities and differences to animal epithelial cells. Open Biol 4, 140017.

4. Dong, J., MacAlister, C.A., and Bergmann, D.C. (2009). BASL controls asymmetric cell division in Arabidopsis. Cell 137, 1320–1330.

5. Rowe, M.H., Dong, J., Weimer, A.K., and Bergmann, D.C. (2019). A Plant-Specific Polarity Module Establishes Cell Fate Asymmetry in the Arabidopsis Stomatal Lineage. bioRxiv, 614636.

6. Houbaert, A., Zhang, C., Tiwari, M., Wang, K., de Marcos Serrano, A., Savatin, D.V., Urs, M.J., Zhiponova, M.K., Gudesblat, G.E., Vanhoutte, I., et al. (2018). POLAR-guided signalling complex assembly and localization drive asymmetric cell division. Nature 563, 574–578.

7. Pillitteri, L.J., Peterson, K.M., Horst, R.J., and Torii, K.U. (2011). Molecular profiling of stomatal meristemoids reveals new component of asymmetric cell division and commonalities among stem cell populations in Arabidopsis. The Plant Cell 23, 3260–3275.

8. Zhang, Y., Wang, P., Shao, W., Zhu, J.K., and Dong, J. (2015). The BASL polarity protein controls a MAPK signaling feedback loop in asymmetric cell division. Developmental cell 33, 136–149.

9. Muroyama, A., Gong, Y., Hartman, K.S., and Bergmann, D. (2022). Cortical polarity ensures its own asymmetric inheritance in the stomatal lineage to pattern the leaf surface. bioRxiv, 2022.2007.2015.500234.

10. Muroyama, A., Gong, Y., and Bergmann, D.C. (2020). Opposing, Polarity-Driven Nuclear Migrations Underpin Asymmetric Divisions to Pattern Arabidopsis Stomata. Current biology : CB 30, 4549–4552.

11. Wallner, E.S., Dolan, L., and Bergmann, D.C. (2023). Arabidopsis stomatal lineage cells establish bipolarity and segregate differential signaling capacity to regulate stem cell potential. Developmental cell 58, 1643–1656.

12. Ruiz Sola, M.A., Coiro, M., Crivelli, S., Zeeman, S.C., Schmidt Kjolner Hansen, S., and Truernit, E. (2017). OCTOPUS-LIKE 2, a novel player in Arabidopsis root and vascular development, reveals a key role for OCTOPUS family genes in root metaphloem sieve tube differentiation. New Phytol 216, 1191–1204.

13. Truernit, E., Bauby, H., Belcram, K., Barthelemy, J., and Palauqui, J.C. (2012). OCTOPUS, a polarly localised membrane-associated protein, regulates phloem differentiation entry in Arabidopsis thaliana. Development (Cambridge, England) 139, 1306–1315.

14. Marhava, P., Bassukas, A.E.L., Zourelidou, M., Kolb, M., Moret, B., Fastner, A., Schulze, W.X., Cattaneo, P., Hammes, U.Z., Schwechheimer, C., et al. (2018). A molecular rheostat adjusts auxin flux to promote root protophloem differentiation. Nature 558, 297–300.

15. Marhava, P., Aliaga Fandino, A.C., Koh, S.W.H., Jelinkova, A., Kolb, M., Janacek, D.P., Breda, A.S., Cattaneo, P., Hammes, U.Z., Petrasek, J., et al. (2020). Plasma Membrane Domain Patterning and Self-Reinforcing Polarity in Arabidopsis. Developmental cell 52, 223–235.e225.

16. Zhang, Y., Bergmann, D.C., and Dong, J. (2016). Fine-scale dissection of the subdomains of polarity protein BASL in stomatal asymmetric cell division. Journal of experimental botany 67, 5093–5103.

17. Breda, A.S., Hazak, O., and Hardtke, C.S. (2017). Phosphosite charge rather than shootward localization determines OCTOPUS activity in root protophloem. Proceedings of the National Academy of Sciences, USA 114, E5721–E5730.

18. Pinak, C., and Devlina, C. (2022). Intrinsically disordered proteins/regions and insight into their biomolecular interactions. Biophysical Chemistry 283, 106769.

19. Yoshida, S., van der Schuren, A., van Dop, M., van Galen, L., Saiga, S., Adibi, M., Moller, B., Ten Hove, C.A., Marhavy, P., Smith, R., et al. (2019). A SOSEKI-based coordinate system interprets global polarity cues in Arabidopsis. Nature plants 5, 160–166.

20. Cao, J., Li, X., and Lv, Y. (2017). Dynein light chain family genes in 15 plant species: Identification, evolution and expression profiles. Plant Sci 254, 70–81.

21. Litchfield, D.W. (2003). Protein kinase CK2: structure, regulation and role in cellular decisions of life and death. Biochem J 369, 1–15.

22. Lucas, J.R., Nadeau, J.A., and Sack, F.D. (2006). Microtubule arrays and Arabidopsis stomatal development. Journal of experimental botany 57, 71–79.

23. Besson, S., and Dumais, J. (2011). Universal rule for the symmetric division of plant cells. Proceedings of the National Academy of Sciences of the United States of America 108, 6294–6299.

24. Muroyama, A., Gong, Y., Hartman, K.S., and Bergmann, D.C. (2023). Cortical polarity ensures its own asymmetric inheritance in the stomatal lineage to pattern the leaf surface. Science 381, 54–59.

25. Ohashi-Ito, K., and Bergmann, D.C. (2006). Arabidopsis FAMA controls the final proliferation/differentiation switch during stomatal development. Plant Cell 18, 2493–2505.

26. Lahav, M., Abu-Abied, M., Belausov, E., Schwartz, A., and Sadot, E. (2004). Microtubules of guard cells are light sensitive. Plant & cell physiology 45, 573–582.

27. Mair, A., Xu, S.L., Branon, T.C., Ting, A.Y., and Bergmann, D.C. (2019). Proximity labeling of protein complexes and cell-type-specific organellar proteomes in Arabidopsis enabled by TurboID. Elife 8, e47864.

28. Branon, T.C., Bosch, J.A., Sanchez, A.D., Udeshi, N.D., Svinkina, T., Carr, S.A., Feldman, J.L., Perrimon, N., and Ting, A.Y. (2018). Efficient proximity labeling in living cells and organisms with TurboID. Nat Biotechnol 36, 880–887.

29. Anne, P., Azzopardi, M., Gissot, L., Beaubiat, S., Hematy, K., and Palauqui, J.C. (2015). OCTOPUS Negatively Regulates BIN2 to Control Phloem Differentiation in Arabidopsis thaliana. Current Biology 25, 2584–2590.

30. Kim, H.S., Park, W., Lee, H.S., Shin, J.H., and Ahn, S.J. (2020). Subcellular Journey of Rare Cold Inducible 2 Protein in Plant Under Stressful Condition. Front Plant Sci 11, 610251.

31. Roeder, A.H., Chickarmane, V., Cunha, A., Obara, B., Manjunath, B.S., and Meyerowitz, E.M. (2010). Variability in the control of cell division underlies sepal epidermal patterning in Arabidopsis thaliana. PLoS biology 8, e1000367.

32. Matos, J.L., Lau, O.S., Hachez, C., Cruz-Ramirez, A., Scheres, B., and Bergmann, D.C. (2014). Irreversible fate commitment in the Arabidopsis stomatal lineage requires a FAMA and RETINOBLASTOMA-RELATED module. Elife 3.

33. Gong, Y., Varnau, R., Wallner, E.S., Acharya, R., Bergmann, D.C., and Cheung, L.S. (2021). Quantitative and dynamic cell polarity tracking in plant cells. New Phytol 230, 867–877.

34. Tyanova, S., Temu, T., and Cox, J. (2016). The MaxQuant computational platform for mass spectrometry-based shotgun proteomics. Nature protocols 11, 2301–2319.

35. Tyanova, S., Temu, T., Sinitcyn, P., Carlson, A., Hein, M.Y., Geiger, T., Mann, M., and Cox, J. (2016). The Perseus computational platform for comprehensive analysis of (prote)omics data. Nat Methods 13, 731–740.

36. Rodriguez-Villalon, A., Gujas, B., Kang, Y.H., Breda, A.S., Cattaneo, P., Depuydt, S., and Hardtke, C.S. (2014). Molecular genetic framework for protophloem formation. Proceedings of the National Academy of Sciences, USA 111, 11551–11556.

37. van Dop, M., Fiedler, M., Mutte, S., de Keijzer, J., Olijslager, L., Albrecht, C., Liao, C.-Y., Janson, M.E., Bienz, M., and Weijers, D. (2020). DIX Domain Polymerization Drives Assembly of Plant Cell Polarity Complexes. Cell 180, 427–439.e412.

38. Willert, K., Brink, M., Wodarz, A., Varmus, H., and Nusse, R. (1997). Casein kinase 2 associates with and phosphorylates dishevelled. The EMBO journal 16, 3089–3096.

39. Halloran, D., Pandit, V., and Nohe, A. (2022). The Role of Protein Kinase CK2 in Development and Disease Progression: A Critical Review. J Dev Biol 10.

40. Pinna, L.A. (2002). Protein kinase CK2: a challenge to canons. J Cell Sci 115, 3873–3878.

41. Jumper, J., Evans, R., Pritzel, A., Green, T., Figurnov, M., Ronneberger, O., Tunyasuvunakool, K., Bates, R., Žídek, A., Potapenko, A., et al. (2021). Highly accurate protein structure prediction with AlphaFold. Nature 596, 583–589.

42. Varadi, M., Anyango, S., Deshpande, M., Nair, S., Natassia, C., Yordanova, G., Yuan, D., Stroe, O., Wood, G., Laydon, A., et al. (2021). AlphaFold Protein Structure Database: massively expanding the structural coverage of protein-sequence space with high-accuracy models. Nucleic acids research 50, D439–D444.

43. Elfmann, C., and Stulke, J. (2023). PAE viewer: a webserver for the interactive visualization of the predicted aligned error for multimer structure predictions and crosslinks. Nucleic acids research 51, W404–W410.

44. Evans, R., O’Neill, M., Pritzel, A., Antropova, N., Senior, A., Green, T., Žídek, A., Bates, R., Blackwell, S., Yim, J., et al. (2022). Protein complex prediction with AlphaFold-Multimer. bioRxiv, 2021.2010.2004.463034.

45. Pace, N.J., and Weerapana, E. (2014). Zinc-binding cysteines: diverse functions and structural motifs. Biomolecules 4, 419–434.

46. Mirdita, M., Schutze, K., Moriwaki, Y., Heo, L., Ovchinnikov, S., and Steinegger, M. (2022). ColabFold: making protein folding accessible to all. Nat Methods 19, 679–682.

47. Barbar, E. (2008). Dynein light chain LC8 is a dimerization hub essential in diverse protein networks. Biochemistry 47, 503–508.

48. Liang, J., Jaffrey, S.R., Guo, W., Snyder, S.H., and Clardy, J. (1999). Structure of the PIN/LC8 dimer with a bound peptide. Nat Struct Biol 6, 735–740.

49. Salinas, P., Fuentes, D., Vidal, E., Jordana, X., Echeverria, M., and Holuigue, L. (2006). An extensive survey of CK2 alpha and beta subunits in Arabidopsis: multiple isoforms exhibit differential subcellular localization. Plant & cell physiology 47, 1295–1308.

50. Moreno-Romero, J., Espunya, M.C., Platara, M., Arino, J., and Martinez, M.C. (2008). A role for protein kinase CK2 in plant development: evidence obtained using a dominant-negative mutant. The Plant journal : for cell and molecular biology 55, 118–130.

51. Meggio, F., and Pinna, L.A. (2003). One-thousand-and-one substrates of protein kinase CK2? FASEB J 17, 349–368.

52. Xu, J., Lee, Y.-R.J., and Liu, B. (2020). Establishment of a mitotic model system by transient expression of the D-type cyclin in differentiated leaf cells of tobacco (Nicotiana benthamiana). New Phytologist 226, 1213–1220.

53. Segui-Simarro, J.M., Austin, J.R., 2nd, White, E.A., and Staehelin, L.A. (2004). Electron tomographic analysis of somatic cell plate formation in meristematic cells of Arabidopsis preserved by high-pressure freezing. Plant Cell 16, 836–856.

54. Canty, J.T., Tan, R., Kusakci, E., Fernandes, J., and Yildiz, A. (2021). Structure and Mechanics of Dynein Motors. Annu Rev Biophys 50, 549–574.

55. Lawrence, C.J., Morris, N.R., Meagher, R.B., and Dawe, R.K. (2001). Dyneins have run their course in plant lineage. Traffic 2, 362–363.

56. Fache, V., Gaillard, J., Van Damme, D., Geelen, D., Neumann, E., Stoppin-Mellet, V., and Vantard, M. (2010). Arabidopsis kinetochore fiber-associated MAP65-4 cross-links microtubules and promotes microtubule bundle elongation. Plant Cell 22, 3804–3815.

57. Liu, A., Mair, A., Matos, J.L., Vollbrecht, M., Xu, S.L., and Bergmann, D.C. (2023). Cell Fate Programming by Transcription Factors and Epigenetic Machinery in Stomatal Development. bioRxiv.

58. Gong, Y., Alassimone, J., Muroyama, A., Amador, G., Varnau, R., Liu, A., and Bergmann, D.C. (2021). The Arabidopsis stomatal polarity protein BASL mediates distinct processes before and after cell division to coordinate cell size and fate asymmetries. Development (Cambridge, England) 148.

59. Zhang, Y., Guo, X., and Dong, J. (2016). Phosphorylation of the Polarity Protein BASL Differentiates Asymmetric Cell Fate through MAPKs and SPCH. Current Biology 26, 2957–2965.

60. Lucas, J., and Geisler, M. (2022). Sequential loss of dynein sequences precedes complete loss in land plants. Plant physiology 189, 1237–1240.

61. Vitre, B., Coquelle, F.M., Heichette, C., Garnier, C., Chretien, D., and Arnal, I. (2008). EB1 regulates microtubule dynamics and tubulin sheet closure in vitro. Nat Cell Biol 10, 415–421.

62. Lin, D., Cao, L., Zhou, Z., Zhu, L., Ehrhardt, D., Yang, Z., and Fu, Y. (2013). Rho GTPase signaling activates microtubule severing to promote microtubule ordering in Arabidopsis. Current biology : CB 23, 290–297.

63. Jespersen, N., Estelle, A., Waugh, N., Davey, N.E., Blikstad, C., Ammon, Y.C., Akhmanova, A., Ivarsson, Y., Hendrix, D.A., and Barbar, E. (2019). Systematic identification of recognition motifs for the hub protein LC8. Life Sci Alliance 2.

64. Strutt, D.I. (2002). The asymmetric subcellular localisation of components of the planar polarity pathway. Semin Cell Dev Biol 13, 225–231.

65. Tree, D.R., Shulman, J.M., Rousset, R., Scott, M.P., Gubb, D., and Axelrod, J.D. (2002). Prickle mediates feedback amplification to generate asymmetric planar cell polarity signaling. Cell 109, 371–381.

66. Radaszkiewicz, K.A., Sulcova, M., Kohoutkova, E., and Harnos, J. (2023). The role of prickle proteins in vertebrate development and pathology. Mol Cell Biochem.

67. Kadrmas, J.L., and Beckerle, M.C. (2004). The LIM domain: from the cytoskeleton to the nucleus. Nat Rev Mol Cell Biol 5, 920–931.

68. Huang, Y., and Winklbauer, R. (2022). Cell cortex regulation by the planar cell polarity protein Prickle1. J Cell Biol 221.

69. Klimczak, L.J., Collinge, M.A., Farini, D., Giuliano, G., Walker, J.C., and Cashmore, A.R. (1995). Reconstitution of Arabidopsis casein kinase II from recombinant subunits and phosphorylation of transcription factor GBF1. Plant Cell 7, 105–115.

70. Bradley, D., and Beltrao, P. (2019). Evolution of protein kinase substrate recognition at the active site. PLoS biology 17, e3000341.

71. Rui, Y., Chen, Y., Kandemir, B., Yi, H., Wang, J.Z., Puri, V.M., and Anderson, C.T. (2018). Balancing Strength and Flexibility: How the Synthesis, Organization, and Modification of Guard Cell Walls Govern Stomatal Development and Dynamics. Front Plant Sci 9, 1202.

72. Paredez, A.R., Somerville, C.R., and Ehrhardt, D.W. (2006). Visualization of cellulose synthase demonstrates functional association with microtubules. Science 312, 1491–1495.

73. Havecker, E.R., Gao, X., and Voytas, D.F. (2005). The Sireviruses, a plant-specific lineage of the Ty1/copia retrotransposons, interact with a family of proteins related to dynein light chain 8. Plant physiology 139, 857–868.

74. Lampropoulos, A., Sutikovic, Z., Wenzl, C., Maegele, I., Lohmann, J.U., and Forner, J. (2013). GreenGate---a novel, versatile, and efficient cloning system for plant transgenesis. PloS one 8, e83043.

75. Bringmann, M., and Bergmann, D.C. (2017). Tissue-wide Mechanical Forces Influence the Polarity of Stomatal Stem Cells in Arabidopsis. Current Biology 27, 877–883.

76. Schindelin, J., Arganda-Carreras, I., Frise, E., Kaynig, V., Longair, M., Pietzsch, T., Preibisch, S., Rueden, C., Saalfeld, S., Schmid, B., et al. (2012). Fiji: an open-source platform for biological-image analysis. Nat Methods 9, 676–682.

77. RStudio Team (2020). RStudio: Integrated Development for R, (Boston, MA: RStudio, PBC.).

78. R Core Team (2022). R: A language and environment for statistical computing, (Vienna, Austria: R Foundation for Statistical Computing).

79. Clough, S.J., and Bent, A.F. (1998). Floral dip: a simplified method for Agrobacterium-mediated transformation of Arabidopsis thaliana. The Plant journal : for cell and molecular biology 16, 735–743.

80. Cox, J., Hein, M.Y., Luber, C.A., Paron, I., Nagaraj, N., and Mann, M. (2014). Accurate proteome-wide label-free quantification by delayed normalization and maximal peptide ratio extraction, termed MaxLFQ. Mol Cell Proteomics 13, 2513–2526.

81. Huala, E., Dickerman, A.W., Garcia-Hernandez, M., Weems, D., Reiser, L., LaFond, F., Hanley, D., Kiphart, D., Zhuang, M., Huang, W., et al. (2001). The Arabidopsis Information Resource (TAIR): a comprehensive database and web-based information retrieval, analysis, and visualization system for a model plant. Nucleic acids research 29, 102–105.

82. Vizcaino, J.A., Cote, R.G., Csordas, A., Dianes, J.A., Fabregat, A., Foster, J.M., Griss, J., Alpi, E., Birim, M., Contell, J., et al. (2013). The PRoteomics IDEntifications (PRIDE) database and associated tools: status in 2013. Nucleic acids research 41, D1063–1069.

83. Elnaggar, A., Heinzinger, M., Dallago, C., Rihawi, G., Wang, Y., Jones, L., Gibbs, T., Feher, T., Angerer, C., Bhowmik, D., et al. (2020). ProtTrans: Towards Cracking the Language of Life’s Code Through Self-Supervised Deep Learning and High Performance Computing. bioRxiv, 2020.2007.2012.199554.

84. Steinegger, M., Meier, M., Mirdita, M., Vohringer, H., Haunsberger, S.J., and Soding, J. (2019). HH-suite3 for fast remote homology detection and deep protein annotation. BMC bioinformatics 20, 473.

85. Vig, J., Madani, A., Varshney, L.R., Xiong, C., Socher, R., and Rajani, N.F. (2020). BERTology Meets Biology: Interpreting Attention in Protein Language Models. bioRxiv, 2020.2006.2026.174417.

86. Fowler, D.M., and Fields, S. (2014). Deep mutational scanning: a new style of protein science. Nat Methods 11, 801–807.

87. Wickham, H. (2016). ggplot2: Elegrant Graphics for Data Analysis, (Springer-Verlag New York).

