## Supplemental figures for "Spatially resolved proteomics of the stomatal lineage: polarity complexes for cell divisions and stomatal pores"

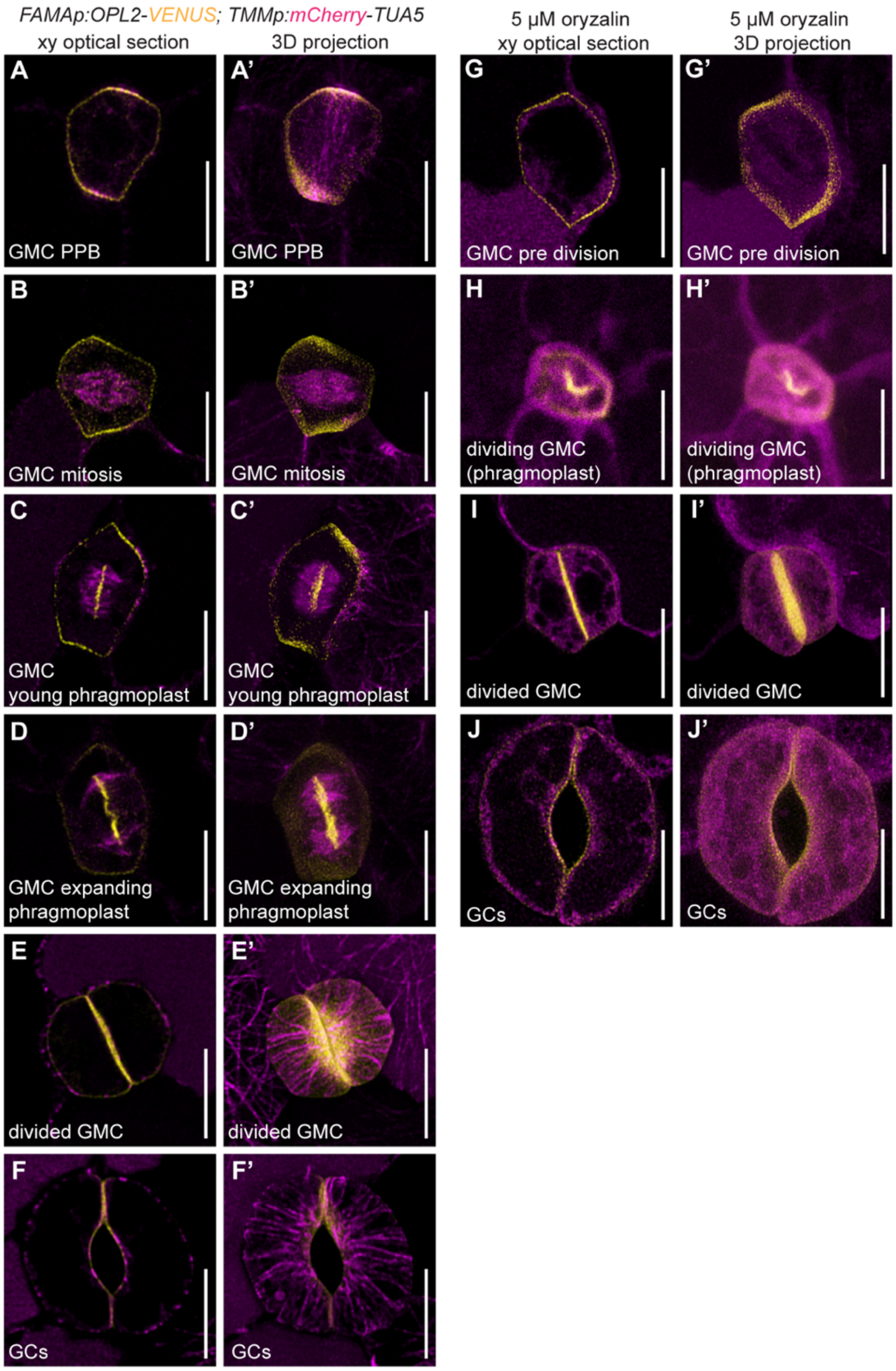


**Figure S1: OPL2 forms polar PM patches before symmetric cell division and attaches to the growing cell plate independently of microtubules (related to Figure 1)**

(**A-F**) GMC and GC specific expression of *FAMAp:OPL2-VENUS* (yellow) and *TMMp:mCherry-TUA5* (magenta) is shown at distinct stages of cell division. The preprophase band (PPB) forms along the long axis of GMCs and runs around the polar patches of OPL2-VENUS (A-A'). During mitosis, polar patches of OPL2-VENUS remain perpendicular to the mitotic spindle polarized to the PM (B-B'). The phragmoplast expands towards these polar patches while OPL2-VENUS also accumulates at the growing cell plate (C-C' and D-D'). OPL2-VENUS occupies the new cell plate after GMC division with cortical microtubules attaching to the ventral wall (E). After pore formation, OPL2-VENUS mostly localizes to contact points of GCs (F-F').

(**G-J**) Treatment with 5 μM oryzalin for 30 min prior to imaging effectively disrupts microtubules at all stages of cell division without affecting OPL2-VENUS polar localization at opposing PM patches (G-G') or localization to the forming ventral wall (H-I) and pore (J), indicating that OPL2 does not require microtubules for polar PM association. Scale bars represent 10 μm.

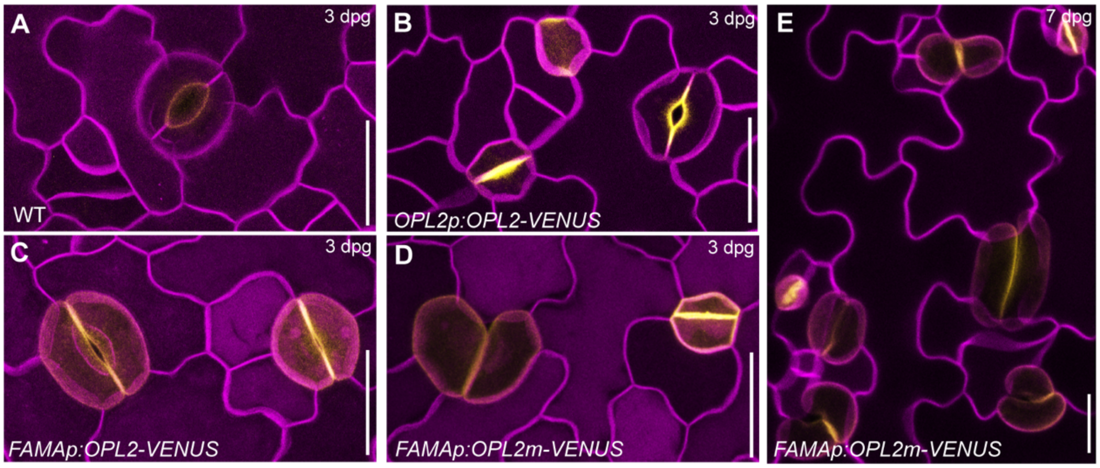


**Figure S2: Manipulation of OPL2 persistence affects stomatal pore development (related to Figure 2)**
**(A-D)** Confocal images of abaxial cotyledons at 3 days post germination (dpg) show a wild-type stoma expressing PM marker *ML1p:mCherry-RCI2A* (magenta signal, A) compared to the same PM marker line expressing *OPL2p:OPL2-VENUS* (yellow). (B) OPL2 in tow patches in young GMCs, along the new division plane as a GMC divides, and marking the pore-forming wall in young GCs. (C-D)GMC and GC-specific *FAMAp* driving expression of wild type OPL2-VENUS (C) or phosphosite mutant OPL2$^{S295K;E296K}$ (OPL2m), a hyperactive OPL protein version [1, 2] that induces deformed, poreless GCs (D). Scale bars represent 20 µm.
**(E)** Pore formation is effectively blocked as GCs expand and distort in *FAMAp:OPL2m-VENUS* expressing lines 7 dpg. Scale bars represent 20 µm.

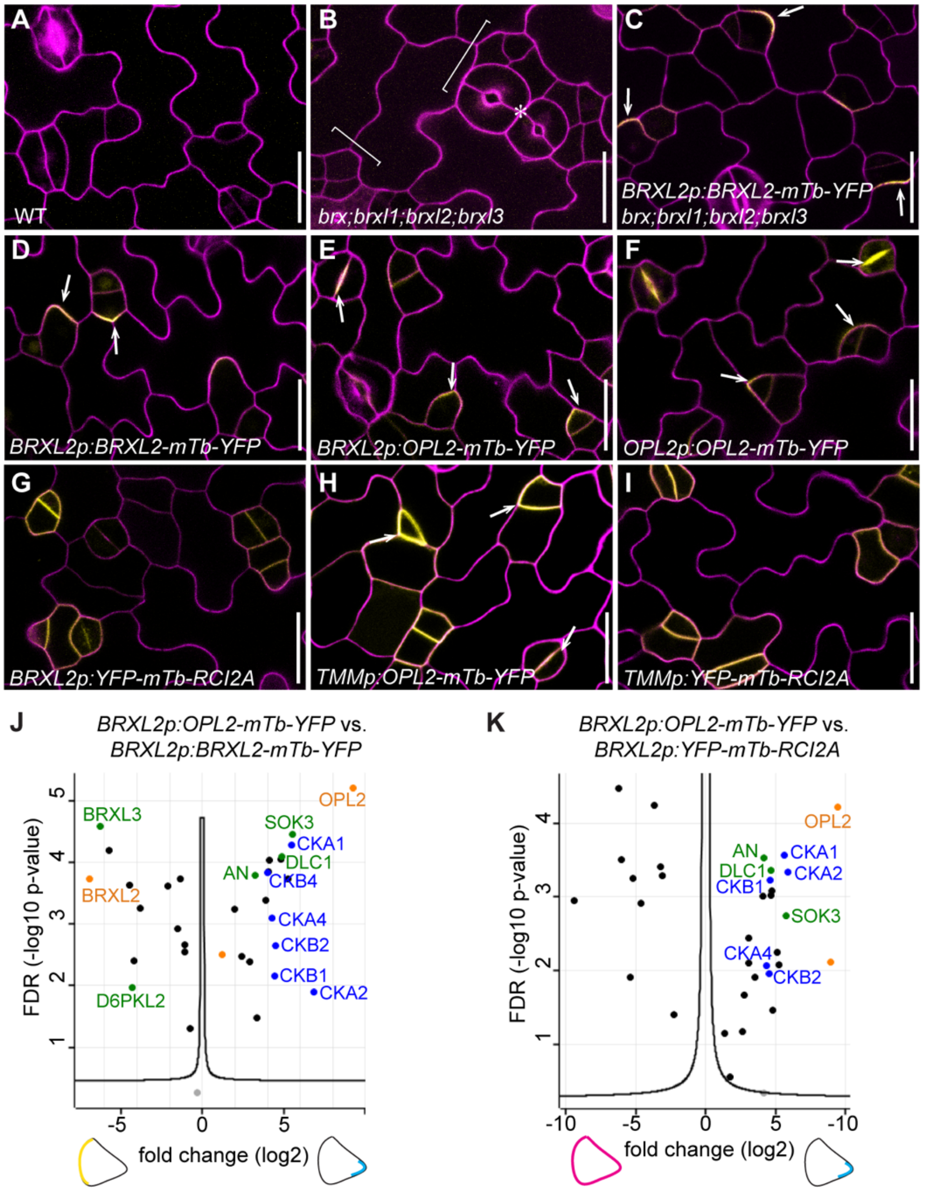


**Figure S3 Proximity labeling lines used in this study with sub-cellular localization of the respective polarity proteins. (related to Figure 3)**

(**A-I**) Confocal images show that *brx;brxl1;brxl2;brxl3* quadruple mutant (*brx-q*) stomatal clustering phenotype (marked by brackets with stomatal pairs marked by asterisk, B) is suppressed by expressing *BRXL2p:BRXL2-mTb-YFP* (C). BRXL2-mTb-YFP polarizes in the *brx-q* background (C) and in wild type (D). *BRXL2p:OPL2-mTb-YFP* (E) and *OPL2p:OPL2-mTb-YFP* (F) and *TMMp:OPL2-mTb-YFP* (H) translational products polarize in ACDs to be inherited to meristemoids and localize at the new cell plate of symmetrically divided GMCs. *BRXL2p:YFP-mTb-RCI2A* (G) and *TMMp:YFP-mTb-RCI2A* (I) fusion products show uniform signals along the PM. Scale bars indicate 20 µm. Arrows point to sites of highest signal intensity.

**(J-K)** Volcano plots depict significantly (FDR = 0.5, S0 = 1) enriched proteins identified between independent sample triplicates expressing *BRXL2p:OPL2-mTb-YFP* vs. *BRXL2p:BRXL2-mTb-YFP* (J) and *BRXL2p:OPL2-mTb-YFP* vs *BRXL2p:YFP-mTb-RCI2A* (K). Independent of the promoter used, OPL2 polarizes in meristemoids and to new cell plates of symmetrically dividing GMCs. Likewise, OPL2-mTb-YFP expressed under the *OPL2, BRXL2* or *TMM* promoter always labels the same top candidate proteins (compare J-K to Figure 2).

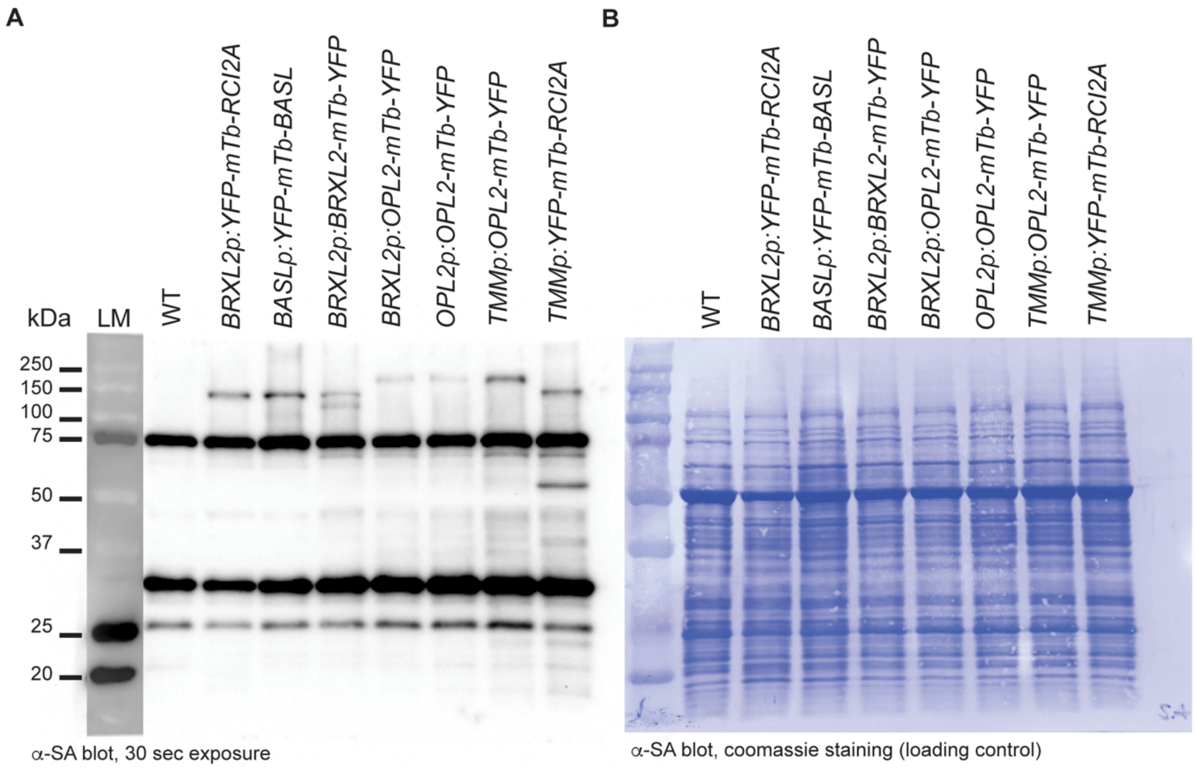


**Figure S4 Western blot analysis of total protein extracts post biotin treatment. (Related to Figure 3)**
(**A**) 100 µl frozen plant powder (collected after 2h treatment with 50 µM biotin) were boiled in 200 µl 4x Laemmli buffer and 5 µl were loaded onto an 10% SDS-PAGE and blotted onto a PVDF membrane, incubated with a streptavidin (SA) antibody before membranes were exposed for 30 sec. WT shows the endogenously biotinylated proteins [3], while all lines expressing a mTb fused to a protein of interest show protein-specific bands, which are especially obvious between 70-150 kDa.
(**B**) Coomassie staining of the blot shown in A serves as a loading control.

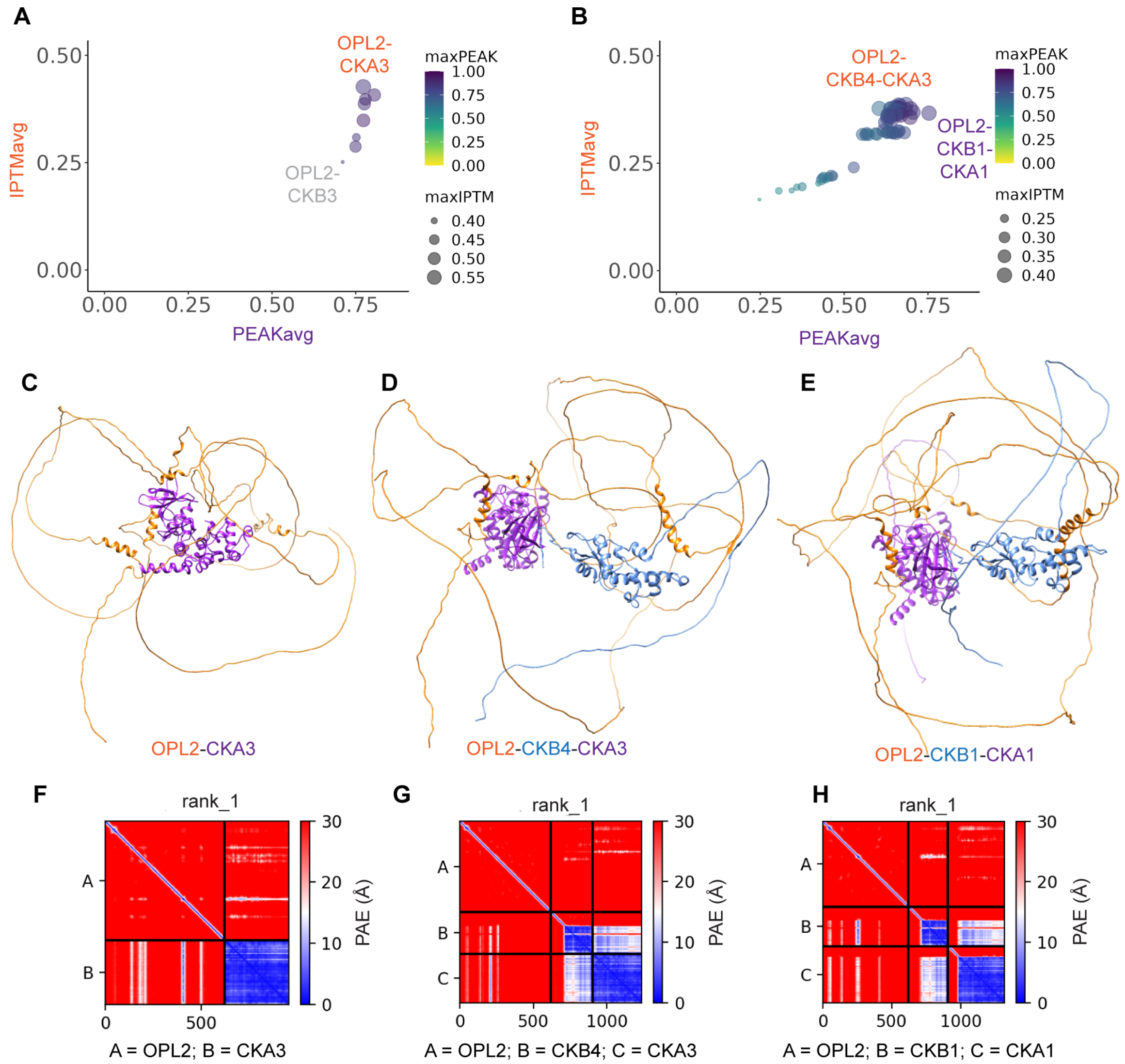


**Figure S5: OPL2 most likely interacts with Casein Kinase II subunits CKA3-CKB4 (Related to Figure 4)**
(**A-E**) AlphaFold-multimer predicted ipTM scores (ipTM: interface predicted Template Modeling score) versus a custom PEAK score representing the inverted and scaled (0-1) minimum of the predicted aligned error (PAE) between the protein chains, excluding intra-molecular interactions. Dimers using OPL2 as bait versus all individual CKII subunits as preys (A) as well as trimers using the OPL2 bait versus all possible CKII subunit combinations (B) were plotted. The highest average ipTM scores were obtained for OPL2- CKA3 (C) and OPL2-CKB4-CKA3 (D) complexes, with CKB4 being a regulatory subunit and CKA3 being the catalytic subunit commonly found in CKII tetramers [4]. The highest PEAK score was obtained for OPL2-CKB1-CKA1 (E).
(**F-H**) Predicted Aligned Error (PAE) plot for OPL2 (chain A) versus CKA3 (chain B, in F) or CKB4 (chain B) and CKA3 (chain C, in G) or CKB1 (chain B) and CKA1 (chain C, in H). Units: amino acid residues; blue to red scale: expected position error in Angstroms.

**References:**


1. Wallner, E.S., Dolan, L., and Bergmann, D.C. (2023). Arabidopsis stomatal lineage cells establish bipolarity and segregate differential signaling capacity to regulate stem cell potential. Developmental cell *58*, 1643-1656.

2. Breda, A.S., Hazak, O., and Hardtke, C.S. (2017). Phosphosite charge rather than shootward localization determines OCTOPUS activity in root protophloem. Proceedings of the National Academy of Sciences, USA *114*, E5721-E5730.

3. Mair, A., Xu, S.L., Branon, T.C., Ting, A.Y., and Bergmann, D.C. (2019). Proximity labeling of protein complexes and cell-type-specific organellar proteomes in Arabidopsis enabled by TurboID. Elife *8*, e47864.

4. Salinas, P., Fuentes, D., Vidal, E., Jordana, X., Echeverria, M., and Holuigue, L. (2006). An extensive survey of CK2 alpha and beta subunits in Arabidopsis: multiple isoforms exhibit differential subcellular localization. Plant & cell physiology *47*, 1295-1308.
